# Effective biotechnology for reducing N_2_O-emissions from farmland: N_2_O-respiring bacteria vectored by organic waste

**DOI:** 10.1101/2023.10.19.563143

**Authors:** Elisabeth G Hiis, Silas H W Vick, Lars Molstad, Kristine Røsdal, Kjell Rune Jonassen, Wilfried Winiwarter, Lars R Bakken

## Abstract

Farmed soils contribute to global warming primarily by N_2_O-emissions, and mitigation has proven difficult. However, a novel approach with promising results in the laboratory, exploits organic wastes both as substrates and vectors for strains of N_2_O-respiring bacteria (NRB), selected for their ability to survive in soil. Here we demonstrate a strong effect in field experiments: fertilization with waste from biogas-production, in which the strain *Cloacibacterium* sp. CB-01 had grown aerobically to ∼6*10^9^ cells mL^-1^, reduced N_2_ O-emissions by 50-95 %. The strong and long-lasting effect of CB-01 is ascribed to it’s tenacity in soil, rather than its biokinetic parameters, which were inferior to other NRB-strains. Scaling up to EU level, we find that national anthropogenic N_2_O-emissions can be reduced by 5-20 %, and more if including other organic wastes. This opens an avenue for cost-effective reduction of N_2_O-emissions for which other mitigation options are currently lacking.

## The nitrogen footprints of farming

Until the mid-20th century, plant production was severely limited by nitrogen, requiring farmers to recycle this element in a reactive form within their agroecosystems. This constraint is reflected in the agricultural treatise by Marcus Porcius Cato (234-143 BC) *De Agri Cultura, which recommends to: “*… save carefully goat, sheep, cattle, and all other dung …” (Hooper and Ash 1934). The invention of the Haber-Bosch process (HB) in 1908 eliminated the nitrogen constraint by producing ammonium from atmospheric nitrogen. HB was a blessing, saving the world from starvation (Erisman et al., 2008), but has become a curse because it allowed farmers to use nitrogen in excess, with marginal penalties for losing nitrogen to the external environment. As a result, agroecosystems have become nitrogen-enriched and leaky, releasing ammonia to the atmosphere and nitrate to the drains, at scales that induce eutrophication and threaten the quality and resilience of both terrestrial and aquatic ecosystems worldwide (Rockström et al. 2009, Sutton et al. 2011, Gu et al. 2023).

Nitrogen fertilisation also causes emissions of the climate gas N_2_O, both from agricultural soils themselves (direct emissions) and from the natural environments due to the input of reactive nitrogen lost from the farms (indirect emissions). These farming-induced emissions account for substantial shares of the escalating concentration of N_2_O in the atmosphere since the industrial revolution (Davidson 2009, Reay et al. 2012, Kanter& Brownlie 2019). A comprehensive analysis of global N_2_O-emissions for 2007-2016 (Tian et al. 2020), estimates that the direct and indirect emissions were 2.3-5.2, and 0.6-2.1 Tg N_2_ O-N y^-1^, in total accounting for > 50 % of the total anthropogenic N_2_O emissions (= 4.1-10.3).

### Mitigation

Reducing the anthropogenic impacts on nitrogen cycling and N_2_O emissions has become a major environmental challenge for the 21st century due to the severity of these issues. An obvious place to start is to improve the nitrogen use efficiency (NUE) of agroecosystems by reducing their losses of ammonia and nitrate (Sutton et al. 2011). This can be achieved by policy instruments to induce shifts in existing-, and implementation of emerging farming technologies (Vatn 2015, Davidson et al. 2014, Zhang et al. 2015, Gu et al. 2023).

While improving NUE can reduce emissions, deliberately manipulating the soil microbes involved holds even greater potential for achieving substantial reductions. Most of the N_2_O emitted from soils is produced by denitrifying bacteria (DB), and denitrifying fungi (DF), and a minor fraction is produced by ammonia-oxidizing archaea (AOA) and -bacteria (AOB) (Bakken and Frostegård 2020). While AOA&AOB and DF are net sources of N_2_O because they lack the enzyme N_2_O-reductase, DB can be both sinks and sources: N_2_O is a free intermediate in their stepwise reduction of nitrate to molecular nitrogen, NO_3_ ^-^ → NO_2_ ^-^ → NO_2_ → N_2_O → N_2_, catalyzed by enzymes encoded by the genes *nar/nap, nirK/S, norB/Q* and *nosZ*, respectively (Zumft 1997). The organisms use this pathway to sustain their respiratory metabolism under hypoxic/anoxic conditions. DB are extremely diverse both regarding their catabolic potential, their regulation of denitrification (Bakken et al. 2012, Lycus et al. 2017), and their denitrification gene sets: a substantial share of DB in soils have truncated denitrification pathways, lacking 1-3 of the 4 genes coding for the complete pathway (Lycus et al. 2017, Pessi et al. 2020). This has been taken to suggest that denitrification is essentially “modular”, i.e. that each step of the pathway is catalyzed by a separate group of organisms, rather than by organisms performing all the steps of the pathway (Roco et al. 2017). The truth is probably a bit of both (Gowda et al. 2021, Hallin et al. 2018).

Being the only sink for N_2_O in soils, the enzyme N_2_O-reductase (NosZ) has been the target for recent attempts to mitigate N_2_O emissions from soils. An intervention that strengthens this sink will lower the N_2_O/N_2_ product ratio of denitrification and hence reduce the propensity of the soil to emit N_2_O into the atmosphere (Bakken and Frostegård 2020, Klimasmith and Kent, 2022). This can be achieved by liming to increase the soil pH: the synthesis of functional nosZ is enhanced by pushing the soil pH towards the upper end of the normal pH-range of farmed soils (pH 5-7) (Liu et al. 2014). As a result, liming acidified soils will reduce their N_2_O emissions by 10-20 %, albeit with a next-to neutral climate effect due to the CO_2_ emission induced by lime application (Henault et al. 2019, Wang et al. 2021).

### Biotechnology for reduced N_2_O emissions

Increasing the soils’ population of N_2_O-respiring bacteria (*NRB*, BOX 1) could decrease the emission of N_2_O (Simon 2021). NRB with a complete denitrification pathway can be net sinks of N_2_O if their denitrification regulatory networks secure earlier and/or stronger expression of NosZ than of the other denitrification enzymes (Lycus et al. 2018, Jonassen et al. 2022a), or if their electron flow is channeled preferentially to NosZ (Gao et al. 2021). Their effect as N_2_O-sinks is plausibly conditional, however, since regulation of their anaerobic respiratory pathway can be influenced by environmental conditions. In contrast, bacteria equipped only with the gene for N_2_O-reduction, will be unconditional net sinks for N_2_O whenever deprived of O_2_ (Shan et al. 2021). In the following, we will call them *Non-denitrifying N_2_O-respiring Bacteria, **NNRB*** because they are unable to denitrify (BOX 1).

We know too little about the ecology and physiology of NNRB to selectively enhance their growth *in situ* (Hallin et al. 2018), but their potential as agents to reduce N_2_O-emissions from soils is indisputable, as demonstrated by laboratory incubations of soils amended with NNRB grown *ex situ* (Domeignoz-Horta et al. 2016). Recently, Jonassen et al. (2022a) suggested that such soil amendment can be done inexpensively on large scale, by using waste from biogas reactors (digestates), destined for soils as organic fertilizers, both as substrate and vector for NRB/NNRB. They successfully enriched and isolated NRB with a strong preference for N_2_O, which could grow aerobically to high cell densities in digestates, and showed that amending soils with NRB-enriched digestates lowered the N_2_O/N_2_ product ratio of denitrification. Their isolates were not ideal, however, because they had genes for the entire denitrification pathway, and their catabolic capacities were streamlined for growth in digestate, not soil. In a follow up study, Jonassen et al. (2022b) designed a dual substrate enrichment strategy, switching between sterilized digestate and soil as substrates, to deliberately select for NRB/NNRB with a broader catabolic capacity and physiochemical tolerance. The enrichments became dominated by strains classified as *Cloacibacterium* (based on 16S amplicon sequencing), and the isolated strain *Cloacibacterium* sp. CB-01 was deemed promising: It carries the genes for NO- and N_2_O-reduction but lacks the genes for reduction of NO_3_ ^-^ and NO_2_^-^, thus qualifying as an NNRB (BOX 1). A subsequent meta-omics analysis of the enrichments and the genome of CB-01 suggested that surface attachment and utilization of complex polysaccharides contributed to its fitness in soil (Vick et al. 2023).

#### BOX 1.

**Bacterial sinks for N_2_O: *NRB* and *NNRB***

Bacteria with a complete denitrification pathway sustain their anaerobic respiration by stepwise reduction of NO_3_ ^-^ to N_2_, catalyzed by four reductases, producing NO_2_ ^-^, NO and N_2_O as free intermediates:

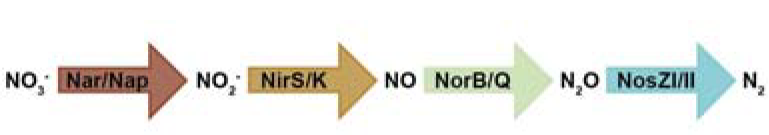

Many bacteria have a truncated pathway, lacking 1-3 of the reductases, with consequences for their role as sources or sinks for N_2_O:

**Terminology:**

***NRB*** = N_2_O-Respiring Bacteria = all bacteria with NosZ. *NRB* equipped with Nir and Nor are either sinks or sources for N_2_O, depending on regulation.

***NNRB*** = Non-denitrifying N_2_O-respiring bacteria = NRB lacking the genes for denitrification *sensus stricto*, i.e. NirK/S. NNRB are net sinks for N_2_O.

**N_2_O-reductase types:** There are two known versions of this copper enzyme: NosZI and NosZII.

Electrons are passed to NosZII via another pathway than NosZI, apparently generating more proton-motive force per electron (Yoon et al. 2016, Hein et al. 2017)

NosZII appears to have a higher affinity for N2O (Yoon et al. 2016).

Here, we have evaluated CB-01’s ability to reduce N_2_O emission from soil, when vectored by digestate. We examined several regulatory and enzyme kinetic traits to assess its inherent strength as an N_2_O sink. Then we tested its capacity in “real life” by conducting field experiments where soils were fertilized with digestate in which CB-01 had been grown to high cell density. Lastly, we assessed the potential of this technology for reducing N_2_O emissions across the European Union.

### Evaluating the respiratory and regulatory phenotype of Cloacibacterium sp. CB-01

The genome of CB-01 contains *nosZII* but lacks any genes coding for dissimilatory reduction of NO_3_ ^-^ and NO_2_ ^-^, predicting a phenotype able to respire N_2_O but neither NO_3_ ^-^ or NO_2_ ^-^, which was confirmed experimentally: in response to oxygen depletion, CB-01 reduced N_2_O to N_2_, but was unable to produce N_2_O from NO_2_ ^-^ (Jonassen et al. 2022b). The fact that it has *norBC*, coding for NO-reductase means that it could produce N_2_O from NO, but the NO kinetics indicates minor NO-reductase activity (Jonassen et al. 2022b Suppl Item 8 B&C). This qualifies CB-01 as an NNRB (see BOX 1), and the laboratory incubation of soils fertilized with digestates containing CB-01 produced marginal amounts of N_2_O (Jonassen et al. 2022b).

The capacity of a strain to reduce N_2_O-emissions is commonly judged by a set of biokinetic parameters (Simon 2021), and we decided to investigate these for CB-01, for comparison with other strains.

#### Growth yield

Based on the bioenergetics and charge separation for aerobic and anaerobic respiration of canonical denitrifying organisms, having NosZI (Box 1), the growth yield in terms of g cell dry weight per mol electrons (*Y*_*e-N2O*_) is ∼60 % of that for aerobic growth (*Y*_*e-O2*_) (van Spanning et al. 2007). For CB-01, which has NosZII, *Y*_*e-N2O*_ was 85 % of *Y*_*e-O2*_ (Fig S1) which lends support to the claim that electron flow via NosZII conserves more energy (by charge separation) than via NosZI (Yoon et al. 2016, Hein et al. 2017).

#### Cell-specific respiration- and growth-rates

Measured aerobic and anaerobic respiration rates during unrestricted growth in nutrient broth at 23 μC were used to estimate maximum growth rates, *μ*_*macx*_, by nonlinear regression (Fig S2), and the maximum rate of electron flow per cell to O_2_ and N_2_O were calculated based on the measured growth yields (*V*_*max*_ = *μ*_*max*_/*Y*). The estimates are *μ*_*max-O2*_ = 0.29 h^-1^ (stdev = 0.006), *μ*_*max-N_2_O*_= 0.11 h^-1^ (stdev = 0.001), *V*_*maxO2*_ = 0.72 fmol O_2_ cell^−1^ h^−1^, *V*_*maxN2O*_= 0.66 fmol N_2_ O cell^-1^ h^-1^. In terms of electron flow rates per cell, we get *V*_*emaxO2*_ = 2.9 fmol e^-^ to O_2_ cell^-1^ h^-1^, *V*_*emaxN2O*_ = 1.3 fmol e^-^ to N_2_O cell^-1^ h^-1^. This shows that CB-01 slows down its respiratory metabolism by ∼50 % when switching from aerobic to anaerobic respiration.

#### Affinity for O_2_ and N_2_O

It is widely accepted that an organism’s ability to effectively mitigate N_2_O emissions is strongly influenced by its affinity to N_2_O. We determined the apparent half saturation constant for O_2_ and N_2_O reduction in CB-01 by nonlinear regression of rates per cell versus concentrations of the two gases in the liquid, and found *k*_*mO2*_ = 0.9 μM O_2_ (SE = 0.27) and *k*_*mN2O*_ = 12.9 μM N_2_O (SE = 1.2) (**Fig S6**). The relatively low *k*_*mO2*_ was expected since the genome of CB-01 contains genes coding for cbb3-type high affinity cytochrome C oxidases (Vick et al. 2023).

#### Comparing the N_2_O sink-strength

To compare CB-01 with other organisms as a sink for N_2_O in soil, we have summarized the biokinetic parameters for various N_2_O-respiring organisms by plotting their “catalytic efficiency” (*V*_*max*_/*K*_*m*_) against their *V*_*max*_ on a cell dry-weight basis (**Fig 1B**) This suggests that CB-01 is far from being the best among N_2_O-respiring organisms: It is on par with the average of others with respect to *V*_*max*_, which is a measure of the N_2_O sink strength at high N_2_O concentrations (>> km=12.9 μM N_2_O ∼ 280 ppmv in the gas phase at 15 ^0^C), while it scores poorly at low N_2_O concentrations (*V*_*max*_*/K*_*m*_ for CB-01 is only 3 % of the average for the others). The apparent bet-hedging (Fig 1A) adds to its inferiority, unless bet-hedging is an “artifact” of the culturing in stirred batches: the fraction of cells expressing NosZ increased with increasing cell density in these batches, and could conceivably reach 1 for cells if clustered in the soil.

**Figure 1.**
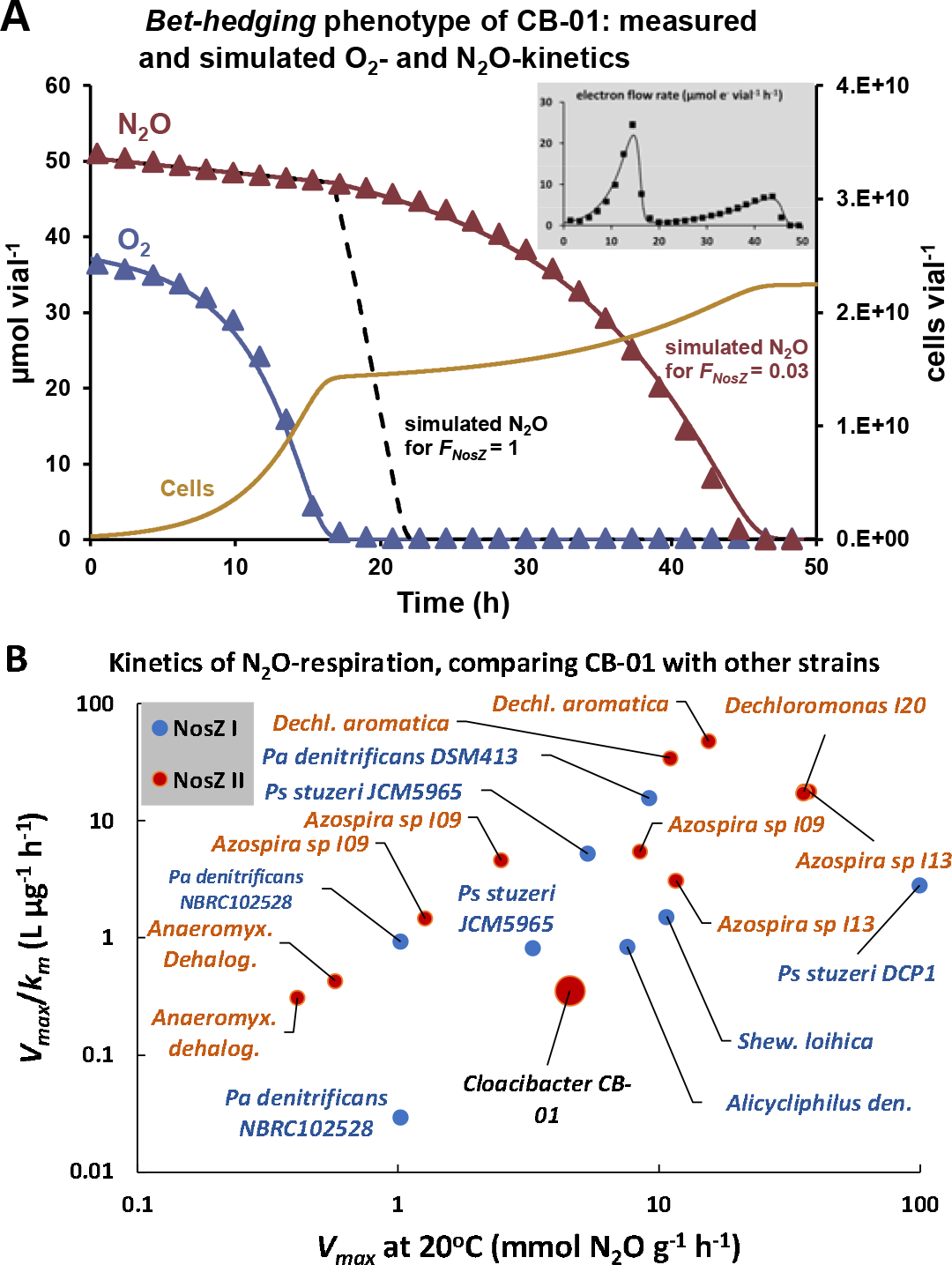
The biokinetics of N_2_O reduction, CB-01 versus other strains. As judged by kinetics of N_2_O respiration in pure culture, CB-01 scores strikingly low compared to other N_2_O-respiring organisms as a sink for N_2_O: the kinetics of N_2_O-respiration in response to O_2_-depletion indicates bet-hedging, i.e. that only a fraction (*F*_*NosZ*_) of the cells express NosZ and start growing by N_2_O-respiration after O_2_-depletion. <u>Panel A</u> illustrates the phenomenon for a single vial: Measured O_2_ and N_2_O (triangles), and simulated (continuous lines) values, using a simplified version of the bet-hedging model of Hassan et al. (2014), with *F*_*NosZ*_ = 0.03. The green line shows the simulated cell density, and the dashed black line shows simulated N_2_O for *F*_*NosZ*_ = 1. The insert shows measured and simulated total electron flow in the vial. <u>Panel B</u> provides a condensed comparison of CB-01 with other N_2_O-respiring organisms regarding its capacity to scavenge N_2_O. Here we have plotted *V*_*max*_*/K*_*m*_ against V (mmol N_2_ O g^−2^ cell dryweight h^−1^) for CB-01 and a range of other organisms with NosZI and NosZII, as measured by others (See Table S1 for details and citations). The comparison shows that CB-01 is close to the average with respect to *V*_*max*_ (37 % of the average of others), while it’s *V*_*max*_ */K*_*m*_ – ratio is very low (∼3 % of the average of others) due to the low apparent affinity for N_2_O (*K*_*m*_ = 13 μM N_2_O).

### The effects of CB-01, vectored by digestate, on N_2_O emissions

CB-01 was found to grow exponentially by aerobic respiration in autoclaved digestate, reaching a cell density of 10^9^ cells mL^-1^ after 20 h. At this point ∼1 % of the organic C in the digestate had been consumed, and growth rate declined gradually, plausibly due to depletion of the most easily available substrate components (**Fig S7**), reaching a final density of ∼6*10^9^ cells mL^-1^ after 2 days, as judged by oxygen consumption and qPCR quantification of CB-01 cells (**Fig S7, Table S2**).

To assess the capacity of such CB-01-enriched digestates to reduce the N_2_O emission from soils under realistic agronomic situations, we conducted three outdoor experiments where N_2_O emissions were measured frequently, using a field flux robot equipped with a tunable diode laser, allowing N_2_O emissions to be measured by 3 minutes enclosures. For details, see **Supplementary 2C**.

The first field experiment (field bucket experiment, described in Supplementary 2D) demonstrated (**Fig 2**) that the initial peak of N_2_O-flux induced by the fertilization with digestate was practically eliminated by CB-01, and that CB-01 continued to have a strong effect throughout: a second peak in N_2_O emission induced by precipitation (day 12) was reduced by 51 %, and the later emission peaks induced by re-fertilization with digestate without CB-01 (indicated by arrows) were reduced by 31, 67, and 46 %.

**Figure 2.**
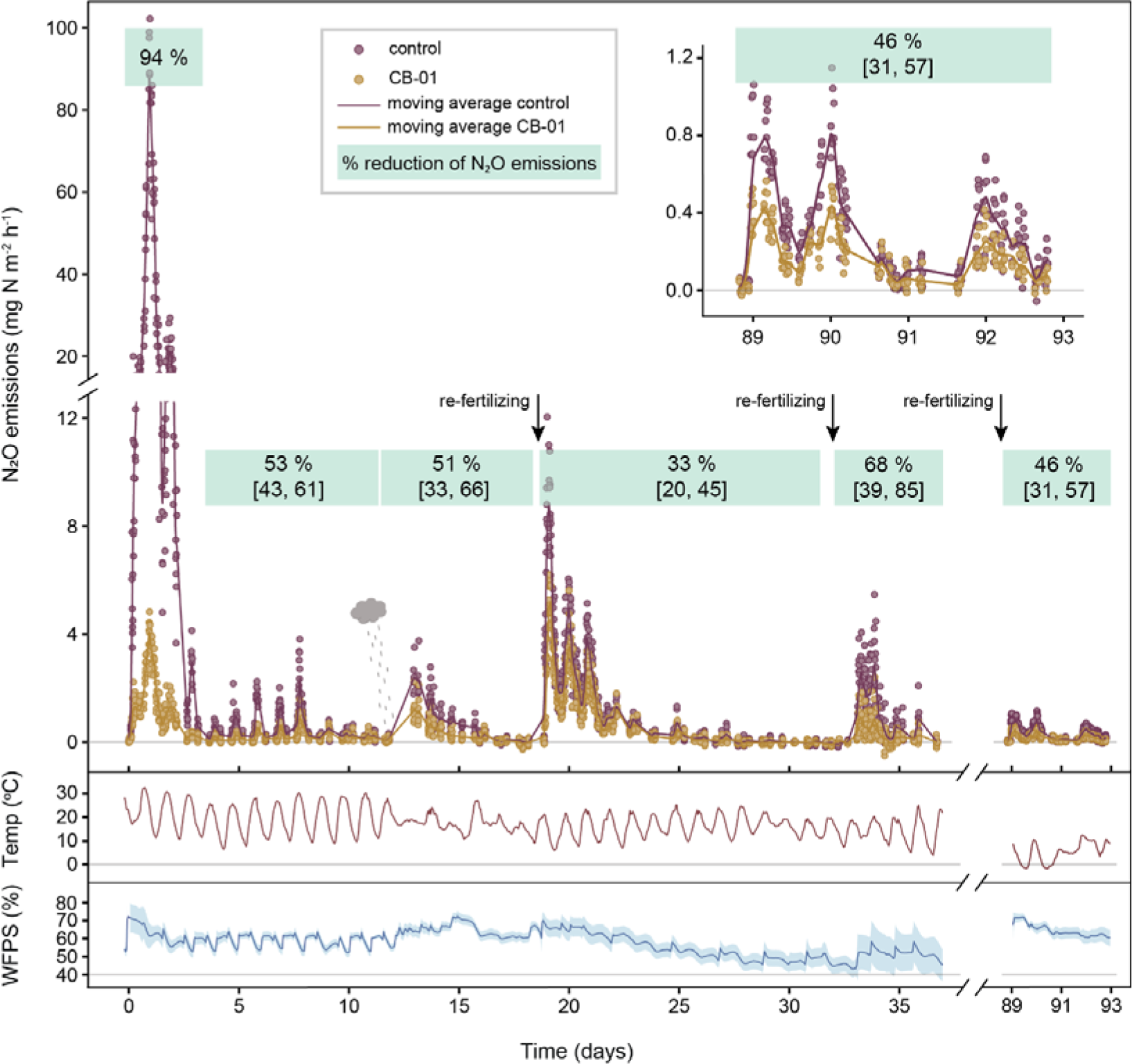
CB-01 effects on N_2_O emission from a clay loam soil pH 6.7. The panel shows N_2_O-flux from buckets with soil throughout 90 days after fertilization (14.07.2021) with digestate (11 L m^−2^) in which the NNRB strain Cloacibacterium sp. CB-01 had been grown to ∼6*10^9^ cells mL^−1^, quantified by qPCR with primers specific for CB-01 (**Supplementary 2F**). Control buckets were fertilized with the same digestate in which CB-01 had been killed by heat (70 °C for 2 hours). The buckets were sown with ryegrass, the soil moisture content was sustained by daily water additions during the first 10days. Buckets were re-fertilized with a lower dose of autoclaved and pH-adjusted digestate without CB-01 (4.6 L m^−2^) after 19, 33 and 89 days. <u>The top pane</u>l shows N_2_O-flux measured by dynamic chamber method (Cowan et al. 2014) with 3 minutes enclosure times, operated by a field robot (**Supplementary 2C**). The emissions are shown as single dots for each enclosure, and with a floating average for each treatment (continuous lines, n = 8 replicate buckets for each treatment), calculated by a Gaussian Kernel smoother). <u>The lower panels</u> show soil temperature (at 0-5.5 cm depth) and water filled pore-space (WFPC). The fluxes show clear diurnal fluctuations, driven by temperature, and transient peaks in response to a rain event (day 12) and in response to re-fertilization (marked by arrows). The % reduction of N_2_O emissions (cumulated flux) by CB-01 were calculated for selected periods, shown in green bars with 95% confidential intervals. The additional control buckets receiving water instead of digestates emitted negligible amounts of N_2_O (result not shown).

Given the number of CB-01 cells added with the digestate (6.6*10^13^ cells m^−2^ soil surface), and the *V*_*max*_ *= 0*.*6* fmol N_2_O cell^−1^ h^−1^ (**Fig S2**), the potential N_2_O-reduction rate, if all the added CB-01 cells were respiring N_2_O at maximum rate, is 1.1 g N_2_O-N m^−2^ h^−1^. The peak N_2_O-flux 1-2 days after fertilization was reduced by ∼85 mg N_2_O-N m^−2^ h^−1^, which is ∼8 % of the estimated potential. For the subsequent peaks of N_2_O-flux, the apparent N_2_O-respiration by CB-01 (i.e. the reduction of the flux) was ≤ 4 mg N_2_O-N m h, which is ≤ 0.36 % of the initial potential. This decline in apparent N_2_O-respiration by CB-01 was plausibly a result of two factors: a gradually declining rate of N_2_O provision by the indigenous microbiome, and a gradually declining number of CB-01 cells.

One would expect that the effect of CB-01 as an N_2_O sink would be marginal in periods with low emissions: Low emissions are due to low waterfilled pore-space (WFPS) (i.e. drained/dry soil), low respiration rate (limited by available organic C-substrates), or both, resulting in marginal hypoxic/anoxic volumes within the soil matrix (Ball, 2013). Under such conditions, the primary source of N_2_O emission could be nitrification (Nadeem et al. 2020), while CB-01 as an N_2_O-sink would be confined to the remaining hypoxic microsites. Inspections of the relationship between the effect of CB-01 and the N_2_O emissions in the control soil (i.e with dead CB-01) lend some support to this: although CB-01 reduced the emissions even for periods with modest emissions, the effect was clearly strongest in periods with high emissions (**Fig 3**).

**Figure 3.**
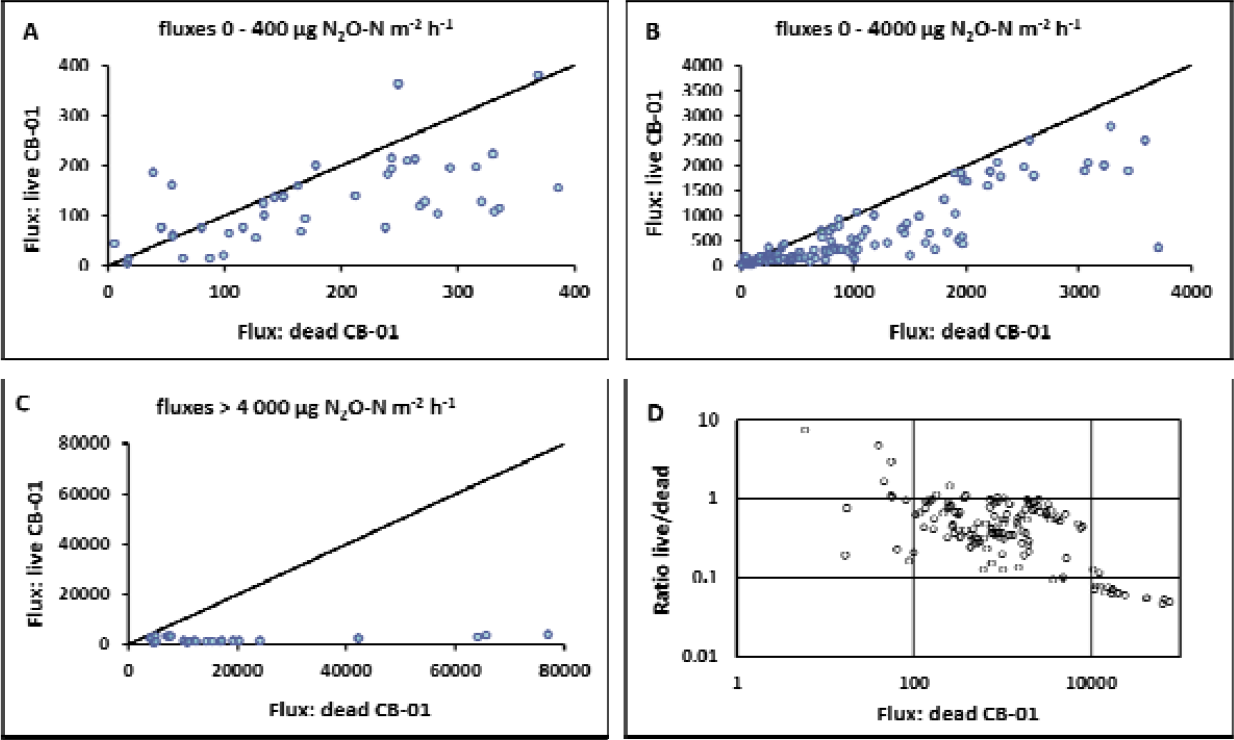
Inspecting the contingent effect of CB-01. Emissions from soil fertilized with digestate containing live *Cloacibacterium* sp. CB-01 plotted against the emissions from soil fertilized with digestate containing dead CB-01 cells (same data as in Figure 2). Panel A-C show the results for the low, intermediate and high emission ranges, respectively. Panel D is a log scaled plot of the ratio between emissions from soil with live and dead CB-01 plotted against the emission from soil fertilized with digestate containing dead CB-01.

We hypothesized that the capacity of CB-01 to reduce N_2_O emissions could be influenced by soil type. Soil pH is plausibly crucial because the synthesis of functional N_2_O-reductase is increasingly impeded by declining pH within the range 4-7, both in CB-01 (Jonassen et al. 2022b) and most other NRB (Bakken and Frostegård 2020). Soil organic carbon content (SOC) could also have an impact. This is because the abundance of CB-01 relative to the abundance of indigenous N_2_O-producing bacteria would be inversely related to SOC, since the abundance of indigenous bacteria in soil is directly related to SOC (Taylor et al. 2002). To explore this, we replicated the bucket experiment (**Fig 2**), but with four different soils spanning a range of pH-levels and including a soil with very high organic carbon content.

The emissions were low compared to the first experiment, plausibly due to lower temperatures (September versus July), however CB-01 significantly reduced the emissions from all four soils. The strong effect in the acidic sandy silt soil (pH = 4.15) was a surprise, until we measured the pH of the soils after amendment with digestate: the incorporation of digestate in this soil increased the pH(CaCl_2_) of the sandy silt soil by more than one pH unit (**Table S3**), reflecting its weak buffer capacity. Most probably, the CB-01 embedded in the digestate experienced even higher pH (pH of the digestate was 7.3). The results for the three clay loam soils show a stronger effect of soil-pH: CB-01 had a clearly stronger effect in the neutral pH clay loam (pH = 6.7) than in the two more acidic clay loams (L: pH = 4.5, O: pH = 5.26).

Finally, we scaled up to a field plot experiment, fertilizing 0.5 m^2^ plots with digestate with live and dead CB-01, carved into the upper 10 cm layer of the soil as in the bucket experiments. The experiment was conducted on field plots that had been limed with 2.3 kg m^-1^ of dolomite in 2014, with an average pH(CaCl_2_) = 6.13 (stdev = 0.10). The high emissions during the first 4 days (Fig 5A) show diurnal variations, peaking when the soil temperature reach their maximum, and a substantial effect of CB-01. Subsequent emissions, measured at low frequency throughout 280 days were much lower, and the effect of CB-01 was not statistically significant, albeit with a wide confidential interval.

**Figure 4.**
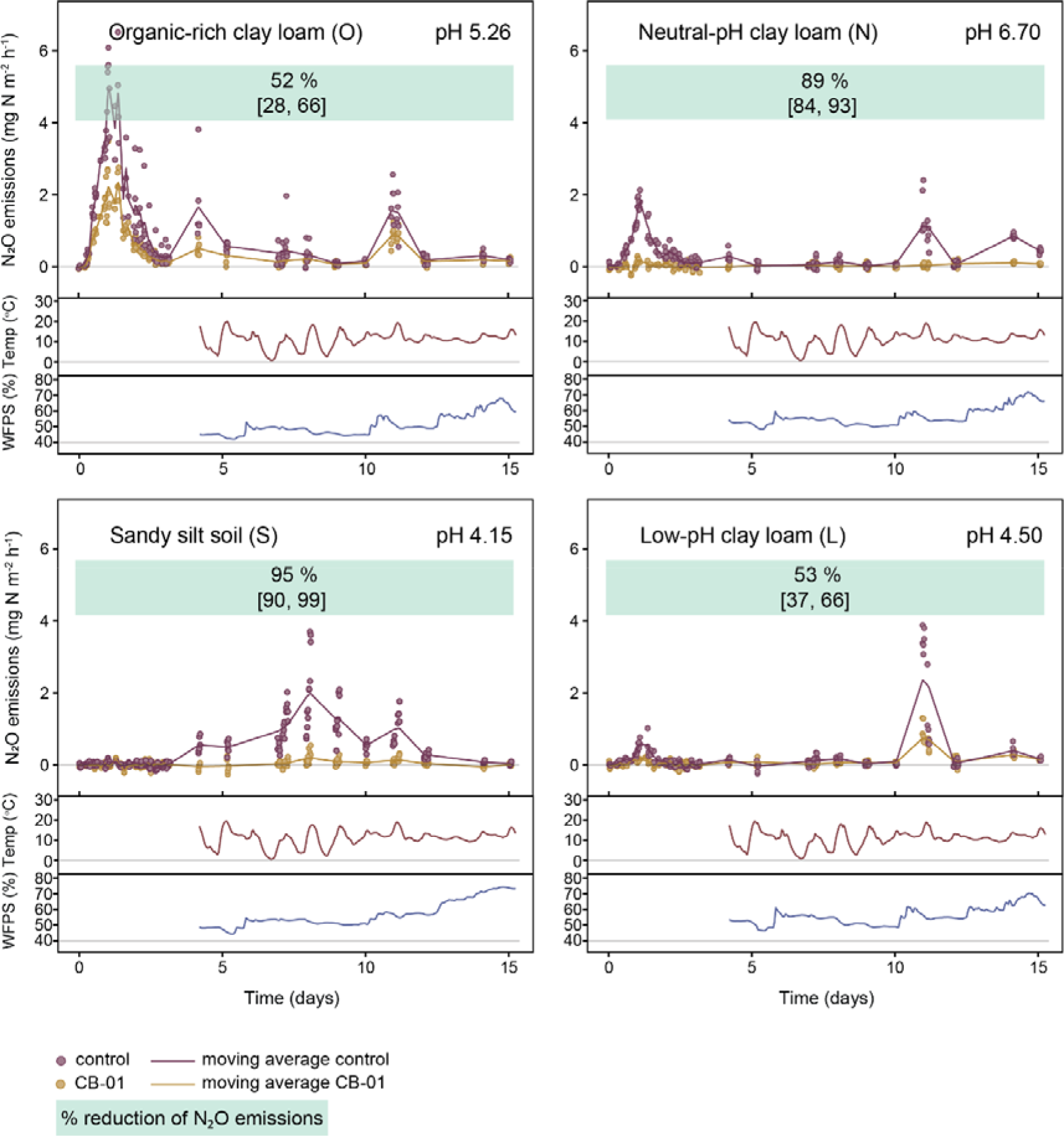
Reduction of N_2_O emissions in different soils in field buckets. The panels show the measured emission after application of digestates with and without CB-01 to four different soils (17.09.2021). The soils’ content of organic C were 15.8 (soil O), 3.21 (soil N), 0.75 (soil S) and 3.23 (soil L) % of dry weight (**Table S2**). The pH(CaCl_2_) prior to fertilization with digestate are given in the panels. The emissions are shown as single dots for each enclosure, and with a floating average for each treatment (continuous lines, n = 6 replicate buckets for each treatment) as in Fig 2. The % reduction of N_2_O emissions (cumulated flux) by CB-01 were calculated for selected periods, shown in green bars with 95% confidential intervals.

**Figure 5.**
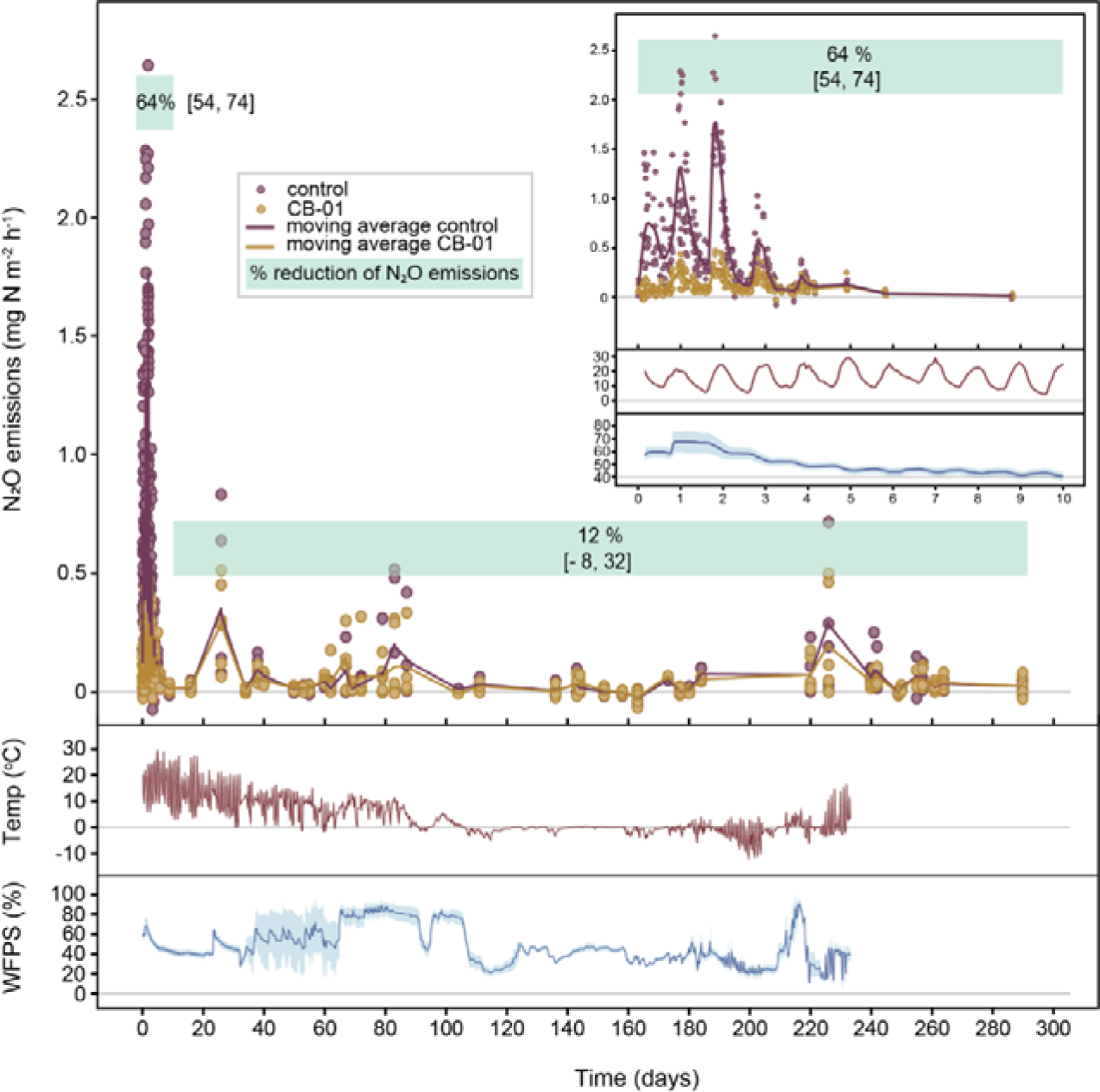
Reduction of N_2_O emissions in field plots. The 0.5 m^−2^ field plots with clay loam pH = 6.13 (soil I, **table S2**) were fertilized by mixing digestate into the upper 10 cm (20.08.2022), with live and dead CB-01 as in previous experiments (n = 6 replicate plots for each treatment). The main panel shows emissions throughout 290 days, and the insert shows emissions during the first 10 days. The % reduction of N_2_O emissions (cumulated flux) by CB-01 for the periods 0-10, and 10-290 days are shown in green bars with 95% confidential intervals.

### Survival of CB-01 in soil, and its effect on the indigenous microbes

Soil microbiome engineering by inoculation/augmentation is an emerging field, promising new possibilities in enhancing agricultural efficiency and sustainability (Lawson et al. 2019). It is challenging, however, because inoculants are invariably found to die out rapidly, plausibly due to a multitude of abiotic and biotic barriers impeding establishment (Kaminsky et al. 2019). CB-01 was obtained through a dual substrate enrichment technique aimed at isolating organisms capable of withstanding the abiotic challenges of soil (Jonassen in 2022b). However, this selection process did not account for the biotic barriers organisms may encounter in soils, such as competition for resources, antagonism, and predation, as highlighted by Albright et al. (2022).

To assess the ability of CB-01 to survive in soil, we used qPCR with specific primers (**Supplementary 2G**) to measure the abundance of CB-01 genomes in soil throughout the long-term field bucket experiment (**Fig 2**), and throughout a laboratory incubation of soil amended with digestate with CB-01 (**Fig 6**). During the laboratory incubation, there was a fast first-order reduction in abundance during day 3-7, and a much slower first order reduction thereafter. In contrast, the abundance was sustained at a high level throughout 90 days in the field buckets, albeit gradually declining. The sustained CB-01 population in the bucket experiment explains why the effect on the N_2_O-emission was sustained (**Fig 2**).

**Figure 6.**
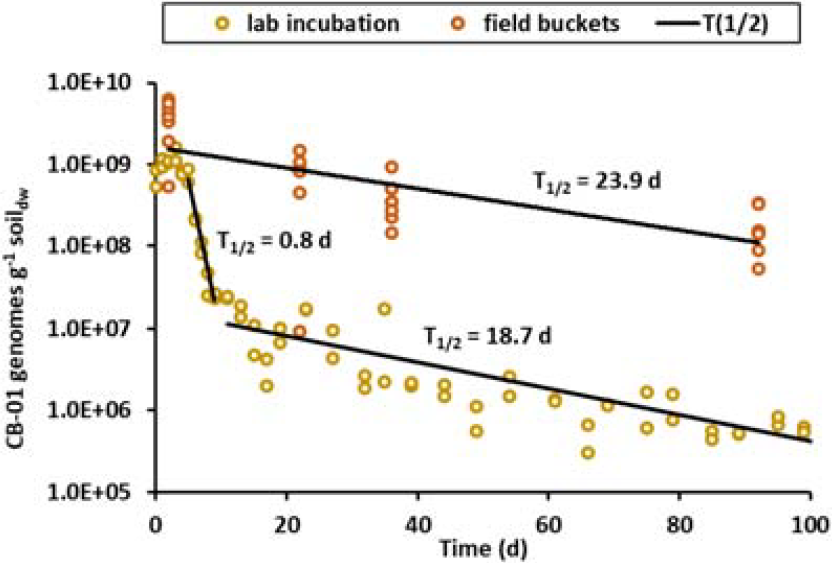
Survival of CB-01 in soil. The abundance of CB-01 was assessed by qPCR as described in Supplementary 2G. The panel shows the genome abundance in the long-term field bucket experiment (Fig 2) and in the laboratory incubation experiment. For the field bucket experiment, where additional digestate was incorporated 2 days before each soil sampling for qPCR. A single dot represents an individual soil sample (n = 8), and the line is the fitted exponential function *N*_*t*_ = *N*_0_*e^*-d*t*^ where *N*_*t*_ is the abundance at time = *t*, and *d* is the apparent first order death-rate (estimated half-life *T*_*1/*2_ = ln(2)/*d*). For the laboratory incubation, three phases can be recognized: an initial apparent growth during the first 2-3 days, followed by a rapid first-order decline during the subsequent 4-5 days, and a slow first order decline thereafter. Of note, the estimated decline in CB-01 genome abundance in the field plot experiment, lasting 280 days indicated very similar average first order death rates (0.028 d^−1^, half life =25 days, Supplementary 4).

But the glaring discrepancy between the field and the lab experiments demands a scrutiny. In the field bucket experiment, digestate (not inoculated with CB-01) were applied three times during the course of the experiment, with soil sampling for quantification of CB-01 abundance conducted two days after each incorporation. Since digestate is a suitable substrate for CB-01, growth of CB-01 in response to each dose could contribute to the sustained population.

Another factor of importance could be that protozoal grazing was plausibly more intense in the laboratory incubations than in the field experiment, due to the higher soil moisture content at the time of CB-01 incorporation: In the lab experiment, the digestate with CB-01 was dripped onto soil which was already very wet (0.53 mL g^−1^ soil dry weight) and retained this high soil moisture throughout. In the field bucket experiment, CB-01-digestate was harrowed into relatively dry soil (0.34 mL g^−1^ soil dry weight), and it remained modestly moist throughout (**Fig 2**). There is ample evidence that low soil moisture protects a bacterial inoculum against protozoal grazing, ascribed to increasing tortuosity, and localization of bacteria in small pores that are inaccessible to the protozoa (van Veen et al. 1997). While we recognize that this is a speculative explanation, it warrants further experimental investigation due to the potential practical implications.

Considering that the soil was heavily inoculated with CB-01 (> 10^9^ cells g^−1^) and that the population was sustained > 10 cells g throughout 90 days in the field buckets, there is a legitimate concern that this could affect the indigenous microbiota (Mallon et al. 2018). We investigated this by analysis of 16S rRNA gene amplicons, excluding the OTU that circumscribed CB-01 (**Supplementary 3**), and found that the digestate as such had a transient impact, but we were unable to discern any consistent difference between the treatments with live versus dead CB-01, which both converged towards the composition of pristine soil (**Fig S12**).

#### Extrapolating national emission reductions

To assess the potential emission reductions by NNRB as compared with other available techniques such as optimized N fertilization and nitrification inhibitors, we estimated emissions for Europe 2030 with the gains model (Winiwarter et al. 2018, Amann et al. 2011).

Consistent with using a uniform emission factor in GAINS (from IPCC, 2006) of 1% of N applied to be emitted as N_2_O, we also assume a uniform factor for emission reductions. From the experiments we conclude that 60% of emission reductions due to NNRB may be considered a conservative estimate. In **Table S5**, emission reductions are shown by EU country for a 2030 scenario if emissions from the application of liquid manure only is reduced by 60%. All other anthropogenic emissions have been left unchanged. Under these assumptions, the total anthropogenic N2O emissions from Europe decrease by 2.7% due to NRB being introduced and applied to all liquid manure systems. This figure is higher in countries that have a high share of liquid manure systems in their agriculture, hence it increases to 4.0% for EU27 (27 EU member countries).

Ongoing work explores the possibility to extend the technology by growing NNRB in all types of organic wastes used to fertilize soils, and by combining the application of mineral N-fertilizers with incorporation of NNRB-amended organic wastes. This requires new strains, technologies and investments, but with a great potential, reducing EU27 agricultural emissions by a third (31%).

It needs to be pointed out that an emission reduction of 60% as derived here for NNRB is much larger than emission reductions typically reported for N2O abatement measures. E.g., GAINS assumes nitrification inhibitors to be able to reduce emissions by as much as 38%, and high-tech mechanical fertilizer-saving technologies (“variable rate application”) to be able to save 24% of the emissions only (Winiwarter et al. 2018).

### Future development

This study presents the first proof of concept, demonstrating a feasible utilization of non-denitrifying N_2_O-respiring bacteria (NNRB) to curb N_2_O emissions from farmland. By using organic waste as substrates and vectors, massive soil inoculation is achieved, which can secure reduced N_2_O-emissions throughout an entire growth season, despite a gradually declining NNRB-abundance. To ensure the robustness and versatility of this biotechnology, we will need an ensemble of new NNRB strains, capable of thriving in waste materials beyond digestates. New NNRB strains will probably vary regarding their ability to tolerate abiotic and biotic stress factors present in the soil. The dual substrate enrichment technique (Jonassen et al. 2022b) selects for strains tolerant of abiotic, but not biotic stress. Consequently, innovative techniques are necessary for selecting strains that tolerate the biotic stress.

## Methods

Determination of the respiratory phenotype was done by batch incubation in the robotized incubation system (**Supplementary 2A**). The same system was used to assess growth of CB-01 in digestate (**Supplementary 2B**), further elaborated by quantifying the abundance by qPCR with specific 16S primers (**Supplementary 2G**).

The effect of CB-01-enriched digestate on N_2_O emission was measured in field bucket experiments (Supplementary 2D) and field plot experiments (**Supplementary 2E**), using a field robot (**Supplementary 2C**). Flux calculations and statistics are described in **Supplementary 2F**. The extrapolation to national emission reductions is described in **Supplementary 2I**.

Survival of CB-01 in the soils in the field bucket experiment was assessed by qPCR with specific 16S primers (**Supplementary 2G**), which was also used to assess the survival in soil incubated in the laboratory (**Supplementary 2H**). The effect of CB-01 on the indigenous microbiota was assessed by analysis of 16S amplicons, general 16S primers (**Supplementary 3**).

