## Supplementary material for "Effective biotechnology for reducing N_2_O-emissions from farmland: N_2_O-respiring bacteria vectored by organic waste"

**­­­**

**Substantial reductions of soil N_2_O emissions utilizing organic waste as vectors for N_2_O-reducing bacteria**

**Supplementary Information**

**Contents:**

**1 The respiratory phenotype of CB-01**

**2 Methods**

**2A Determining the phenotype by robotized batch-cultivation**

**2B Culturing CB-01 in digestate for fertilization experiment**

**2C Monitoring N_2_O emission by a field robot**

**2D Field-bucket experiments**

**2E Field plot experiments**

**2F Calculations of emissions and statistical analyses**

**2G Tracing CB-01 in digestate and soil**

**2H Survival of CB-01 in soil, laboratory incubation**

**2J** **Extraplolating to national emission reductions**

**3 Effect of CB-01 on the soil microbiome**

**4 Survival of CB-01 in the field plot experiments**

**1** **The respiratory phenotype of CB-01**

Non-denitrifying N_2_O-respiring bacteria (NNRB) have attracted much interest recently as net sinks for N_2_O in soils, potentially curbing N_2_O emissions if abundant (Hallin et al. 2018, Simon 2021). NNRB-strains vary grossly in their apparent capacity to act as N_2_O-sinks, assessed by determining their biokinetic parameters: NNRB strains are commonly assumed to be strong N_2_O sinks if they have strong affinity (low apparent *k_m_*) for N_2_O and a high maximal rate of N_2_O reduction (*V_max_*), or simply a high catalytic efficiency, i.e. a high V_max_/k_m_ (Yoon et al. 2016). Another desirable, albeit speculative feature would be to reduce N_2_O under aerobic or at least hypoxic conditions (Kim et al. 2022).

To assess *Cloacibacterium* sp. CB-01 along these criteria, and to compare it with other strains, we conducted an in-depth investigation of its respiratory phenotype using a robotized incubation system (Molstad et al. 2016) which provides high resolution gas kinetics (CH_4_, CO_2_, O_2_, NO, N_2_O and N_2_) in batch cultures in gas tight vials, as they deplete oxygen and switch to anaerobic respiration, reducing N_2_O to N_2_. Combined with adequate calculation of gas transport, this approach has proven powerful in unravelling novel regulatory features such as *bet-hedging* (Hassan et al. 2014, Lycus et al. 2018), as well as characterizing key enzyme parameters *in vivo* (Hassan et al. 2016a).

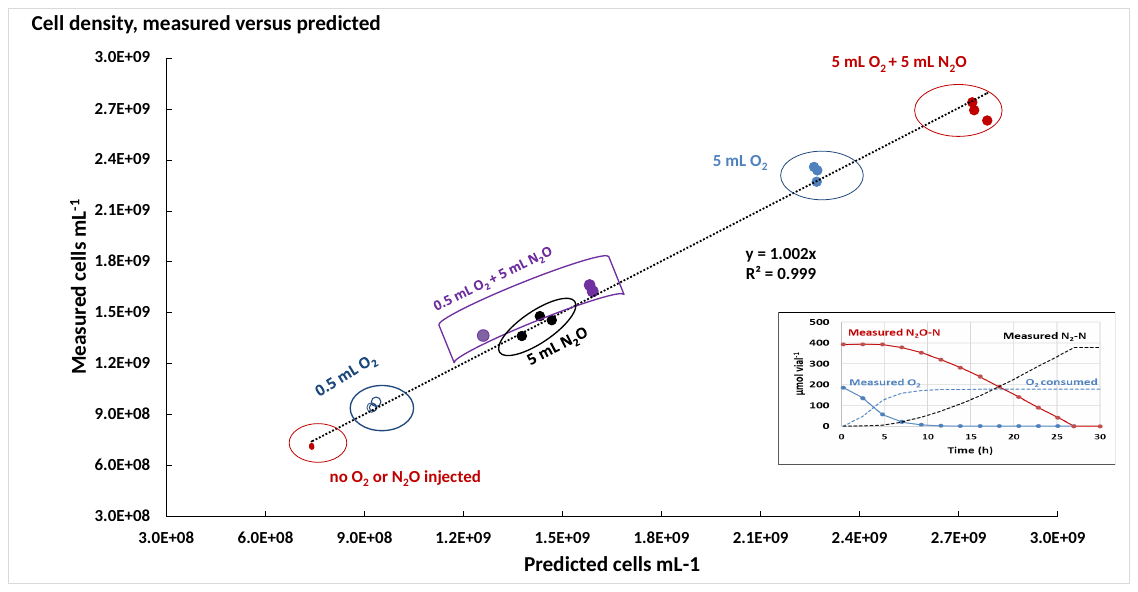

**Fig S1: Growth yield by aerobic and anaerobic respiration.** The growth yield of CB-01 by aerobic and anaerobic respiration was assessed by batch cultivation in 50 mL nutrient broth in gas tight 120 mL serum vials, stirred by magnetic bars. Prior to inoculation, the 18 vials were He-washed by repeated evacuation and He-filling (Molstad et al. 2007) and provided with different amounts of N_2_O and O_2_ by injection with a syringe. They were then placed in the thermostatic water-bath (temperature = 23 ^o^C) of the robotized incubation system (Molstad et al. 2016). After temperature equilibration and subsequent release of overpressure due to N_2_O- and O_2_-injection, the vials were inoculated with 3.5*10^10^ cells (7*10^8^ cells mL^-1^). The incubation system monitors the O_2_, N_2_O and N_2_ concentration in the headspace, and these measurements were used to estimate the cumulated reduction of O_2_ and N_2_O throughout the incubation for each vial. The inserted panel shows an example for a single vial with 5 mL N_2_O and 5 mL O_2_ injected. The cumulated O_2_- and N_2_O-reduction (to N_2_) do not add up to 100 % of the initial amounts because each sampling removes a fraction of the headspace gas (replaced by helium, see Molstad et al. 2007).

When all the electron acceptors were depleted, the cell density was measured by OD at 600 nm. The relationship between cell density and OD_600_ was measured in separate experiments with suspensions of CB-01 with a range of densities (microscopic counts), which showed a linear relationship for OD_600_ ≤ 0.5 (cell density = 3.34*10^9^ mL^-1^ OD^-1^). Thus, any sample with OD_600_>0.5 was diluted to reach OD_600_<0.5 for determination of cell density. In the same experiment, the cell dry weight of CB-01 was determined by weighing dry cells (cells washed three times in distilled water by dispersion and centrifugation, then dried at 105^o^C)). The dry weight was 108 fg cell^-1^ ± SE= 7.5 (n=9).

The measured yield per mol of N_2_O and O_2_ was found by using the Generalized Reduced Gradient Solver in Excel for the entire dataset. The panel shows the result for individual vials, as a plot of the predicted cell density (based on the yields given below) against measured cell density. The estimated yields were Y_N2O_= 1.7*10^14^ cells mol^-1^ N_2_O and Y_O2_=4*10^14^ cells mol^-1^ O^2^. The yields per mol electrons are Y_e-N2O_ = 0.85*10^14^ mol^-1^ e^-^ to N_2_O and Y_e-O2_= 1*10^14^ cells mol^-1^ e^-^ to O_2_. The yields in terms of dry weight g (given the dry weight= 108 (± SE= 7.5) fg cell^-1^), are **Y_e-N2O_ = 9.2 ± 0.6 g mol^-1^ e^-^ to N_2_O and Y_e-O2_ = 11 ± 0.8 g mol^-1^ e^-^ to O_2_.**

In comparison, Bergaust et al. (2010) found *Paracoccus denitrificans* to have Y_e_-_O2_ = 3.75*10^13^ cells mol^-1^ e^-^ to O_2_ = 11.2 g cell dry weight mol^-1^ e^-^ to O_2_ (cell dry weight=298 fg), which is practically identical with Y_e-O2_ determined for CB-01. Y_e-N2O_ was 85% of Y_e-O2_ ratio for CB-01, which is surprisingly high compared to that measured for *P. denitrificans* (53%), and compared to the expectations (~0.6) based on the charge separation per electron for aerobic and anaerobic respiration for denitrifying organisms carrying NosZ clade I (van Spanning et al. 2007). However, there is mounting evidence that some organisms with NosZ Clade II have higher yields per electron to N_2_O than organisms with NosZ clade I (Yoon et al. 2016), suggesting that the electron pathway to NosZ Clade II generates more charge separations than the pathway to NosZ Clade I, which is thermodynamically possible (Hein and Simon 2019, Simon 2021).

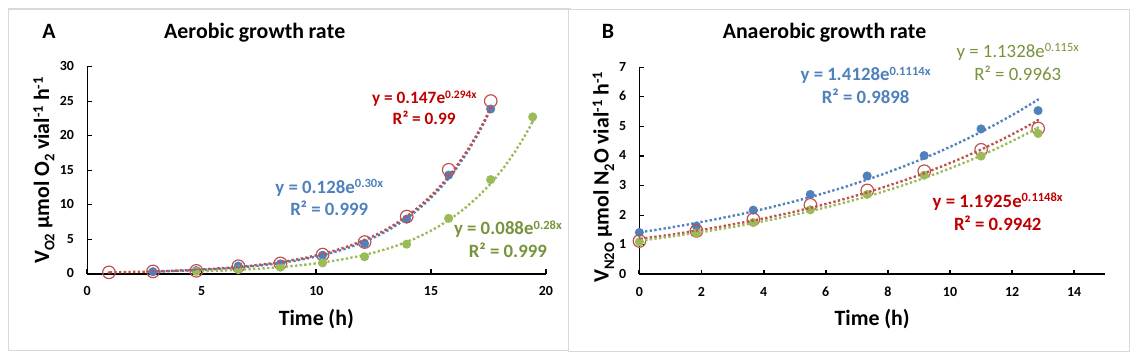

**Fig S2. Aerobic and anaerobic growth rates**. The aerobic and anaerobic growth rates were determined by nonlinear regression of the rates of O_2_- and N_2_O-reduction against time during unrestricted growth in nutrient broth in stirred batch cultures in 120 mL serum vials (23 ^o^C). Panel A shows the rate of oxygen consumption in three replicate vials with 6 vol% O_2_ in the headspace, inoculated with ~7*10^8^ cells mL^-1^. The oxygen concentration in the liquid was 80 µM initially, declining to 50 µM at the end, thus growth was not restricted by the availability of O_2_. The growth rate in each vial was estimated by nonlinear regression (equations and R^2^ values shown in the panel) and the average aerobic growth rate is 0.29 h^-1^ (stdev = 0.006). Panel B shows the rate of N_2_O reduction in three replicate vials after O_2_-depletion, when the cell density had reached ~1.7*10^10^ vial^-1^. The N_2_O concentration in the liquid was 280 µM initially, declining to 110 µM at the end (14 h), thus growth was not restricted by availability of N_2_O. The anaerobic growth rate was calculated for each vial as in panel A, and the average is 0.11 h^-1^ (stdev = 0.001) Given these aerobic and anaerobic unrestricted growth rates, we can calculate V_max_ per cell for O_2_ and N_2_O (*V_max_* = µ/Y). The estimated values are *V_maxN2O_*= 0.6 fmol N_2_O cell^-1^ h^-1^, *V_maxO2_* = 0.72 fmol O_2_ cell^-1^h^-1^. The dry weight of the CB-01-cells is 108 fg, thus V_maxN2O_ expressed on a dry weight basis is 0.0059 mol N_2_O g^-1^ dry weight h^-1^. The maximal electron transport rates for aerobic and anaerobic respiration are *V_emaxO2_* = 2.9 fmol e^-^ to O_2_ cell^-1^ h^-1^, *V_max e N2O_*= 1.2 fmol e^-^ to N_2_O cell^-1^ h^-1^. Thus, the aerobic respiratory rate of CB-01 is more than twice the anaerobic respiration rate.

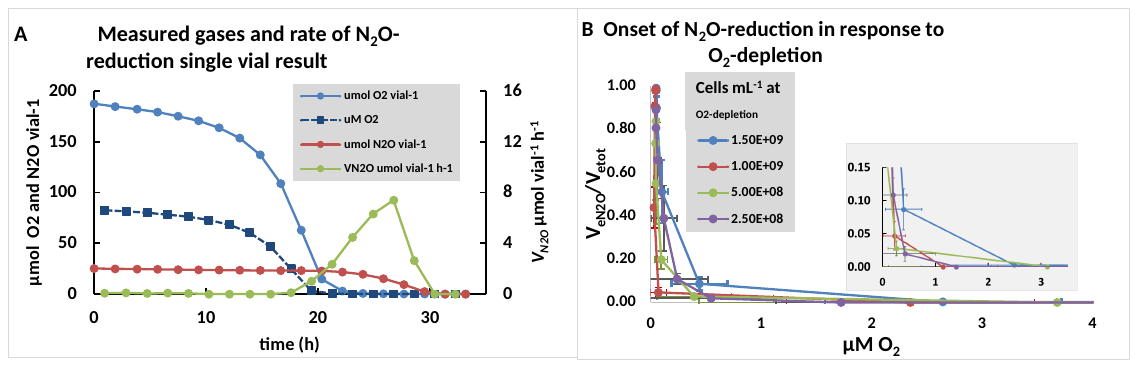
**Fig S3 Onset of N_2_O-reduction during O_2_ depletion.** Denitrifying bacteria vary as to how early they initiate anaerobic respiration during O_2_-depletion (Bergaust et al. 2011). To explore this for *Cloacibacterium* sp. CB-01, we ran several experiments, both in nutrient broth (Panel A-C) and in digestate (panel D). Panel A&B: show the results for experiments where the inoculum was raised through >10 generations under strict aerobic conditions, thus diluting out any N_2_O reductase that might be present in the cells. Panel A shows the gas measurements, the O_2_-concentration in the liquid as calculated from the O_2_ transport rate (see Molstad et al. 2007), and the rate of N_2_O-reduction (*V_N2O_*). Panel B shows the ratio *V_eN2O_*/*V_etot_*, i.e. the fraction of total electron flow that goes to N_2_O (for each time increment), plotted against the O_2_ concentration in the liquid. Panel C shows V_eN2O_/V_etot_ (plotted against [O_2_]) for an experiment where the inoculum had been exposed to hypoxia, thus with NosZ expressed already. Panel D shows the result for growth in digestate, inoculated with cells raised aerobically.

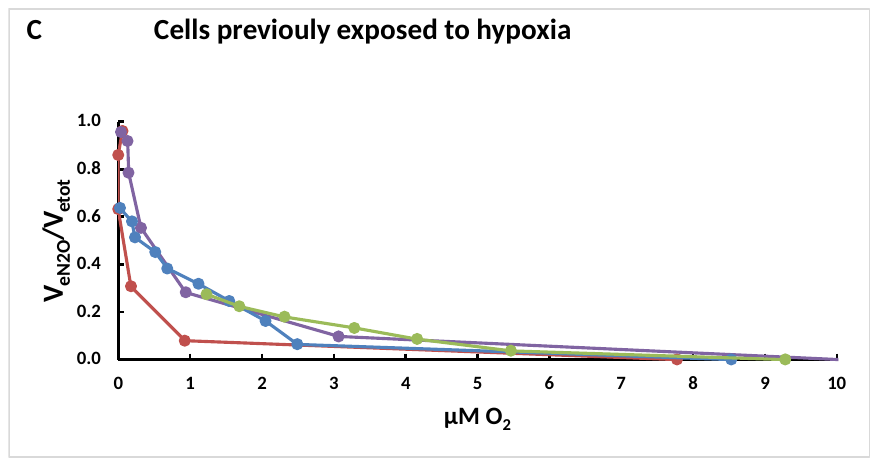

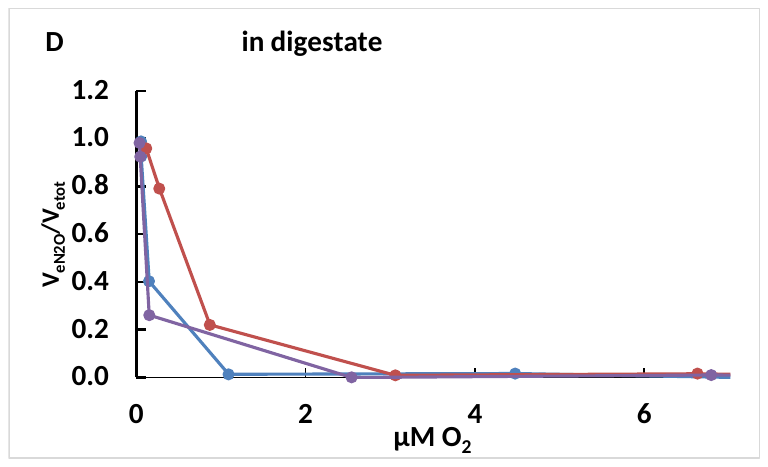

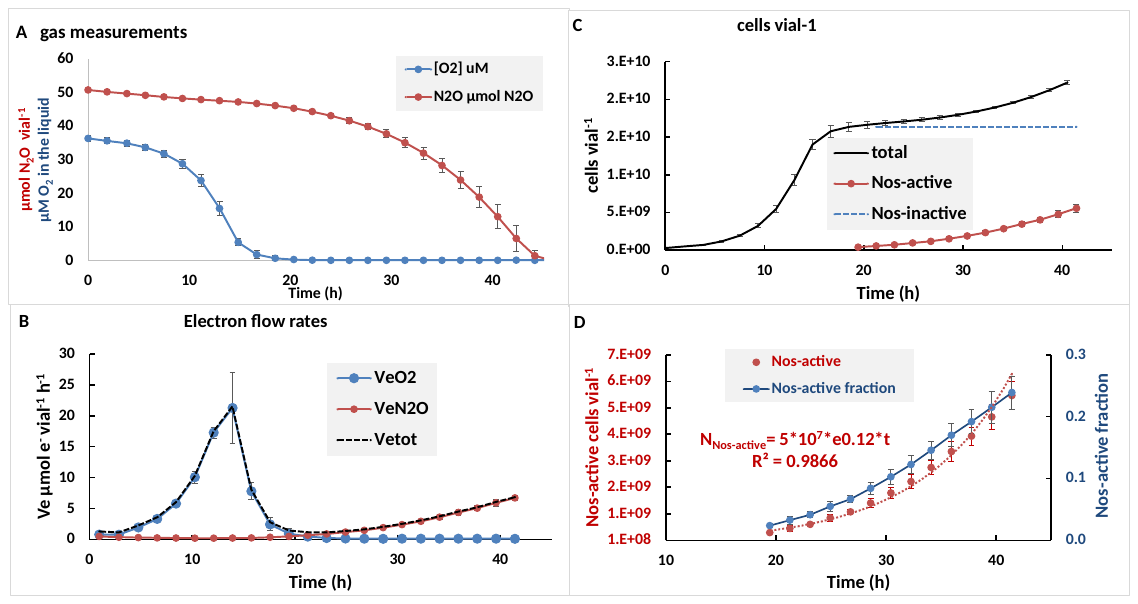

**Fig S4** **The electron flow kinetics during transition from aerobic to anaerobic respiration suggests *bet-hedging*.** To investigate the characteristic denitrification regulatory phenotype of CB-01, 120 mL vials with 50 mL nutrient broth and O_2_ + N_2_O in He-atmosphere were inoculated with 2.7*10^8^ cells per vial, which had been raised under strict aerobic conditions (n=3 replicate vials). The vials were monitored for gas kinetics as the cultures grew by oxygen initially, and then switched to respiring N_2_O in response to O_2_-depletion. The experiment included four treatments, all with 1 mL N_2_O (50 µmol N_2_O vial^-1^), but five different amounts of O_2_ (0, 0.8, 1.4, 3 and 4.6 mL O_2_ (3 replicate vials for each O_2_ level). The panel shows the result for the vials with 0.8 mL. Panel A shows the measured amounts of O_2_ and N_2_O per vial. Panel B shows the electron flow rates to O_2_ and N_2_O (and total electron flow rate as a dashed line) as calculated from the measured O_2_ and N_2_O. Two phenomena stand out here: as oxygen was depleted, the electron flow rate declined to very low values and the subsequent electron flow rate to N_2_O increased exponentially, with an apparent growth rate of 0.12 h^-1^ (which is slightly higher than the anaerobic growth rate of CB-01 determined previously). This is the typical pattern for a *bet-hedging* denitrifying organism, i.e. an organism which expresses denitrification enzymes only in a fraction of the cells (Hassan et al. 2014, Lycus et al. 2017). Assuming this, we investigated the possible fraction of cells that express NosZ and engaged in anaerobic respiration and growth: Panel C shows the estimated total number of cells (based on the cumulated O_2_ and N_2_O-consumption and the yields per mol O_2_ and N_2_O, **Fig S1**), and the number of N_2_O-respiring cells (Nos-active) calculated from the measured N_2_O-reduction rate (*V_N2O_* , mol N_2_O vial^-1^ h^-1^) and the assumption that V_maxN2O_ = 0.65 fmol N_2_O cell^-1^ h^-1^ (as determined previously): NosZ-active cells vial^-1^= V_N2O_/*V_maxN2O_*. The blue dashed line is the estimated number of cells without Nos (Nos inactive), assumed to be cells entrapped in anoxia without Nos, hence unable to synthesize Nos. Panel D shows the number of Nos-active cells as numbers per vial, and as fraction of the total number of cells in the vial. This fraction increases with time due to growth by N_2_O-respiration, and the fraction at the time of O_2_ depletion is a crude estimate of the fraction of cells which were able to express Nos before O_2_ is completely exhausted.

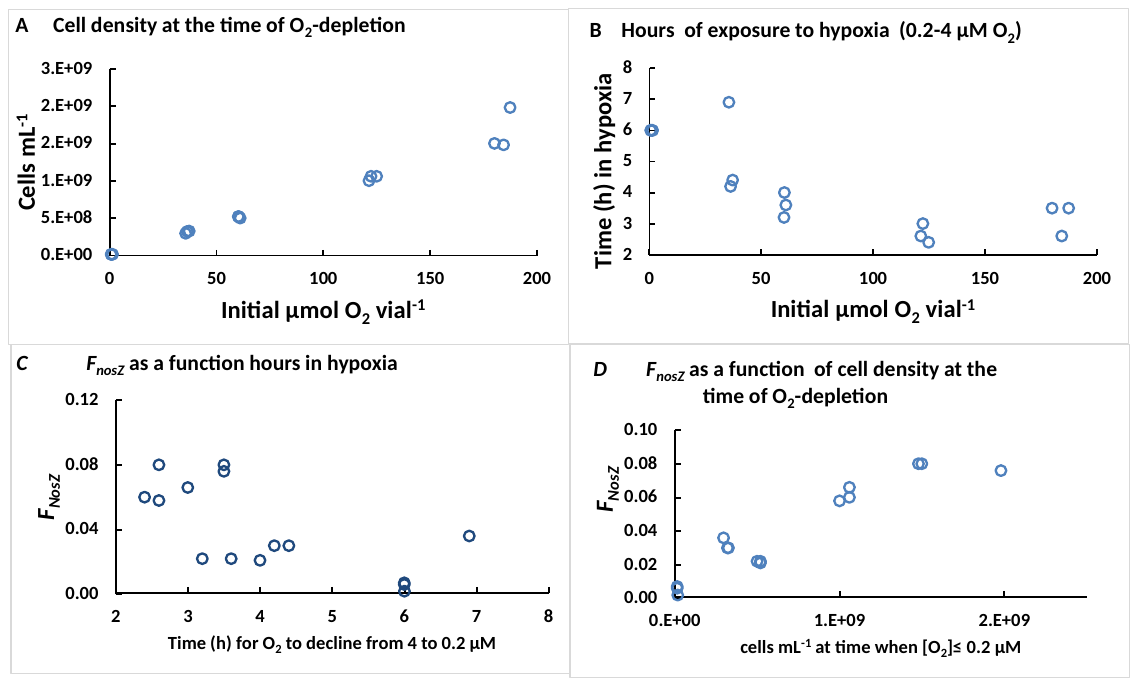
 **Fig S5 *Bet-hedging* depending on cell density at the time of O_2_ depletion.** Lycus et al. (2018) provided experimental proof for *bet-hedging* in *Paracoccus denitrificans*, hypothesized by Hassan et al. (2014) based on modelling of the type of electron flow kinetics throughout the transition from oxic to anoxic respiration as shown in **Fig S4**. In *P. denitrificans*, the fraction of cells that were able to switch to anaerobic respiration (***F_den_***), thus avoiding entrapment in anoxia (Kellermann et al. 2022), was proportional to the time length of the hypoxic phase preceding complete anoxia. This was ascribed to a stochastic initiation of transcription (of *nirS*, coding for nitrite reductase) once the cells experience hypoxia. To investigate if the apparent *bet-hedging* in CB-01 shows the same pattern, we estimated ***F_nosZ_*** for all the 15 vials in the experiment reported in **Fig S4**: the vials were all provided with 1 mL N_2_O but five different amounts of O_2_ (0, 0.8, 1.4, 3 and 4.6 mL O_2_, n= 3 replicate vials for each O_2_-level). The cell density at the time of O_2_-depletion increased with increasing initial O_2_ (Panel A), while the time length of exposure to hypoxia (arbitrarily defined as 0.2-4 µM O_2_) declined (Panel B). A simplified version of the *bet-hedging* model (Hassan et al. 2016b), assuming instantaneous expression of NosZ in a fraction of the cells (***F_NosZ_***) as O_2_ reached below 0.5 µM, was fitted to observed gas kinetics (O_2_ and N_2_O) for each vial to estimate ***F_NosZ_*** (all other parameters were as determined previously (**Fig S1-3**). Contrary to our expectations, the estimated fraction of cells expressing nosZ (***F_NosZ_***) decreased with increasing time length of exposure to hypoxia (Panel C) and increased with cell density at the time of O_2_ depletion (Panel D). Interestingly, practically all cells appeared to become entrapped in anoxia (***F_den_*** = 0.002-0.006) in the vials without any O_2_ injected. The results warrant further investigations to provide direct evidence for the cell differentiation (*bet-hedging*), and the mechanism causing ***F_den_*** to increase with cell density. A tantalizing hypothesis is that quorum sensing induction of NosZ expression is involved.

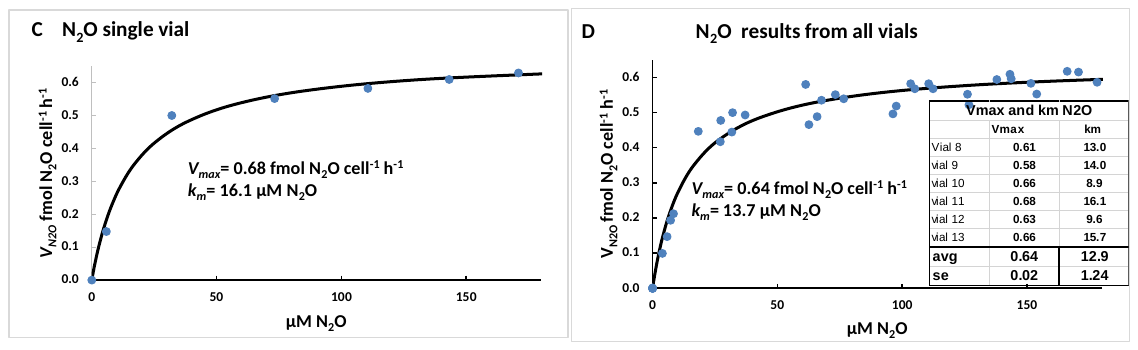

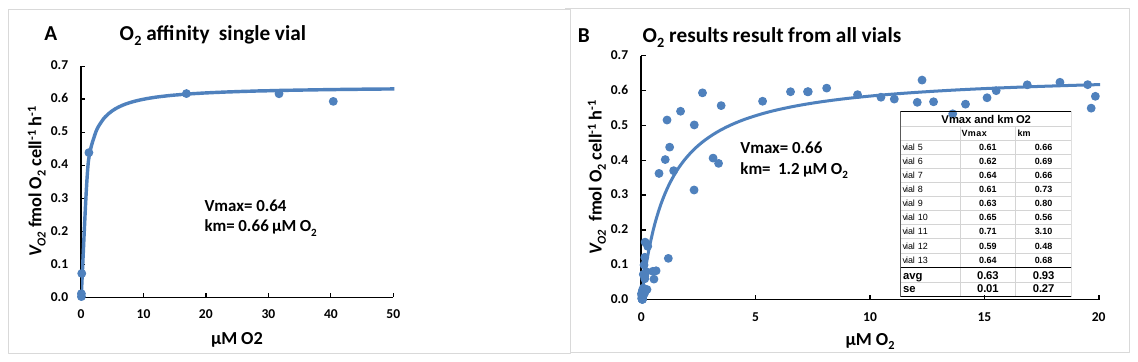
**Fig S6. Apparent affinity for O_2_ and N_2_O.** To assess the affinity for O_2_ and N_2_O, we measured the rates of O_2_- and N_2_O-reduction as the batch cultures depleted the two electron acceptors. The rates as measured (mol vial^-1^ h^-1^) were converted to rates per actively respiring cell (*V_O2_* and *V_N2O_*, fmol cell^-1^ h^-1^) based on the numbers of active cells in the vial at each time point (= the midpoint between two samplings). The number of active cells were the numbers of cells in the inoculum + new-grown cells as calculated from the cumulated consumption of O_2_ and N_2_O (and the growth yield per mol, **Fig S1**), as done previously for determining the affinity for NO (Hassan et al. 2016a). For *V_N2O_*, the number of actively N_2_O-respiring cells was only a fraction of the total (as shown in **Fig S5** panels C&D). The maximum rates V_max_ and the apparent *k_m_* values were found by fitting the Michaelis Menten model V=*V_max_**S/(*k_m_*+S) to the data (S is the concentration of O_2_ and N_2_O in the liquid) by least square, using the Generalized Reduced Gradient Solver in Excel. This was done for each individual vial, and for the collective datasets. Panels A and B show the results for O_2_, for a single vial (A) and for the entire dataset (B). Embedded in the panel is the estimated *k_m_* for each individual vial. Panels C and D show the results for N_2_O, for a single vial (A) and for the entire dataset (B). Embedded in the panel is the estimated *k_m_* for each individual vial. These results show a relatively strong affinity for O_2_ (*k_m_*~1 µM O_2_), and rather weak affinity for N_2_O (*k_m_*~13 µM N_2_O).

**Table S1. Comparison of biokinetic parameters for CB-01 and other N_2_O-respiring bacteria**. The maximum rates of N_2_O-respiration (V_max_), and the affinity for N_2_O, normally expressed as the half saturation concentration (*K_m_*), have been measured in various N_2_O-respiring strains to assess and compare their capacity to scavenge N_2_O in soil. A plausible way to rank strains according to their capacity to reduce N_2_O emission at low N_2_O-concentrations is to calculate their *V_max_*/*K_m_* ratios, since this approximates the slope of the rate against N_2_O-concentration ([N_2_O]) at very low concentrations ([N_2_O]<<*K_m_*). At very high N_2_O-concentrations, the capacity to reduce emissions is proportional with *V_max_*. In the table below, we have listed *V_max_* and *K_m_* determined for a range of N_2_O-respiring strains. Some report *V_max_* as fmol N_2_O cell^-1^ h^-1^, while others report it as mol g^-1^ cell dry weight h^-1^. To enable a comparison, we have converted all *V_max_* values to mol N_2_O g^-1^ cell dry weight h^-1^, assuming a cell volume = ~0.6 µm^3^ and 200 fg dry weight per cell (Bakken and Olsen 1983), and converted all values to V_max_ at 20 ^0^C assuming the rates increase exponentially with temperature by a factor of 2.5 per 10 ^0^C increase in temperature (Q_10_=2.5). The calculated *V_max20_/K_m_* values are plotted against *V_maxN20_* in Figure 1 (Panel B) in the main paper.

|  | **nosZ** | **temp** | **Vmax** | **Vmax** | **Km** | **V_max20_** | **V_max20_/K_m_** | **Reference** |
| --- | --- | --- | --- | --- | --- | --- | --- | --- |
| **Organism** | **Clade** | **(^o^C)** | **fmol**  **cell^-1^ h^-1^** | **Mol g^-1^ DW h^-1^** | **µM N_2_O** | **mmol g^-1^ h^-1^**  **at 20^O^C** | **L µg^-1^ h^-1^**  **at 20^0^C** |  |
| *Pseudomonas stutzeri* DCP1 | I | 30 |  | 0.250 | 35.5 | 99.840 | 2.812 | Yoon et al. 2016 |
| *Shewanella loihica* PV4 | I | 30 |  | 0.027 | 7.07 | 10.704 | 1.514 | Yoon et al. 2016 |
| *Paracoccus denitrificans* DSM413 | I | 20 |  | 0.027 | 0.59 | 26.760 | 45.356 | Hassan et al. 2016b |
| *Pseudomonas stutzeri* JCM5965 | I | 30 | 1.64 | 0.008 | 4.01 | 3.280 | 0.818 | Suenaga et al. 2018 |
| *Paracoccus denitrificans* NBRC102528 | I | 30 | 0.51 | 0.003 | 34.8 | 1.020 | 0.029 | Suenaga et al. 2018 |
| *Paracoccus denitrificans* NBRC102528 | I | 30 | 0.51 | 0.003 | 1.1 | 1.020 | 0.927 | Qi et al. 2022 |
| *Pseudomonas stutzeri* JCM5965 | I | 30 | 2.66 | 0.013 | 1.01 | 5.320 | 5.267 | Qi et al. 2022 |
| *Alicycliphilus denitrificans* I51 | I | 30 | 3.78 | 0.019 | 8.98 | 7.560 | 0.842 | Suenaga et al. 2019 |
| *Dechloromonas aromatica* RCB | II | 30 |  | 0.028 | 0.324 | 11.064 | 34.148 | Yoon et al. 2016 |
| *Anaeromyxobacter dehalogenans* 2CPC | II | 30 |  | 0.001 | 1.34 | 0.410 | 0.306 | Yoon et al. 2016 |
| *Cloacibacterium* sp. CB-01 | II | 23 |  | 0.006 | 12.9 | 4.558 | 0.353 | this study |
| *Azospira* sp. I09 | II | 30 | 0.634 | 0.003 | 0.868 | 1.268 | 1.461 | Suenaga et al. 2018 |
| *Azospira* sp. I13 | II | 30 | 5.8 | 0.029 | 3.76 | 11.600 | 3.085 | Suenaga et al. 2018 |
| *Azospira* sp. I09 | II | 30 | 1.24 | 0.006 | 0.54 | 2.480 | 4.593 | Qi et al. 2022 |
| *Azospira* sp. I13 | II | 30 | 18.84 | 0.094 | 2.12 | 37.680 | 17.774 | Qi et al. 2022 |
| *Dechloromonas* sp. I20 | II | 30 | 18 | 0.090 | 2.04 | 36.000 | 17.647 | Suenaga et al. 2019 |
| *Azospira* sp. I09 | II | 30 | 4.23 | 0.021 | 1.55 | 8.460 | 5.458 | Suenaga et al. 2019 |
| *Azospira* sp. I13 | II | 30 | 17.9 | 0.090 | 2.1 | 35.800 | 17.048 | Suenaga et al. 2019 |
| *Dechloromonas aromatica* RCB | II | 30 | 7.748 | 0.039 | 0.324 | 15.496 | 47.827 | Suenaga et al. 2019 |
| *Anaeromyxobacter dehalogenans* 2CP-C | II | 30 | 0.287 | 0.001 | 1.34 | 0.574 | 0.428 | Suenaga et al. 2019 |

**2 Methods**

**2A	Robotized batch cultivations for determination of respiratory phenotypes**

The respiratory phenotypic parameters of CB-01 were determined by batch culturing in the robotized incubation system designed and described by Molstad et al. (2007, 2016). The system hosts up to 30 parallel stirred batch cultures (normally 50 mL) in 120 mL gas tight serum vials with He-atmosphere (with or without N_2_O and O_2_), which are sampled frequently for measuring the concentrations of O_2_, N_2_, N_2_O, NO and CO_2_. Robust routines are established for calculating the rates of production/consumption of all the gases (taking sampling-loss and leakage into account), and for calculating gas concentrations in the liquid as a function of gas concentrations and the rate of transport between liquid and headspace. These routines are included in a spreadsheet which is publically available, including a set of instruction videos (Bakken 2021). The system has been used in numerous investigations of the respiratory phenotypes of denitrifying bacteria (Bergaust et al. 2008, 2010; Hassan et al. 2016ab, Jonassen et al. 2022ab, Gao et al. 2021, Kellerman 2022, Qu et al. 2016).

To enable refined analyses of the respiratory phenotype of CB-01, we initially determined the cell dry weight (fg cell^-1^), and the growth yields for aerobic (*Y_O2_*, cells mol^-1^ O_2_) and anaerobic (*Y_N2O_*, cells mol^-1^ N_2_O) respiration by measuring the cell yields in batches provided with various amounts of O_2_ and N_2_O. This enabled inspection of the cell specific respiration rates (fmol cell^-1^ h^-1^) throughout subsequent batch incubations, based on measured rates (mol O_2_ & N_2_O vial^-1^ h^-1^) for each time interval between two gas samplings, and the estimated cell number in the vial for the same time interval (=N_ini_ + *Y_O2_**CumO_2_+*Y_N2O_**CumN_2_O, where N_ini_ is the initial number of cells at time=0, CumO_2_ and CumN_2_O are the cumulated consumption of the two gases). The cell specific rates calculated this way allowed an analysis of the affinity for O_2_ and N_2_O by plotting cell specific rates of O_2_ and N_2_O against the concentrations of the two gases in the liquid as the cultures depleted the gases, and fitting the Michaelis-Menton function to these data (least square). Batch cultures provided with both N_2_O and O_2_ in the headspace were monitored as they depleted O_2_ and switched to respiring N_2_O, thus determining the critical concentration of O_2_ (in the liquid) at which the cells started to respire N_2_O. The kinetics of electron flow throughout such transitions from aerobic to anaerobic respiration were used to assess the fraction of cells expressing N_2_O-reductase in response to O_2_-depletion, using a simplified version of the model developed by Hassan et al. (2016b).

**2B	Culturing CB-01 in digestate for field experiments**

For each field experiment, fresh digestate was collected from a wastewater treatment plant close to Oslo (VEAS), described in Jonassen et al. (2022a). Averaged values of the quality parameters for the period of digestate collection were: dry matter content = 3.97 weight % (stdev=0.16), ignition loss of dry matter = 55.6 % (stdev=2), pH = 7.72 (stdev=0.07) and NH_3_+NH_4_^+^ = 1.71 g N L^-1^ (stdev=0.12).

Prior to cultivation of CB-01*,* the digestate was heat-treated, aerated and pH-adjusted. For the field bucket experiments (Chapter 2B), the digestate was autoclaved (121 °C for 20 min), and then sparged with air (while stirred) for 48 hours to secure chemical oxidation of Fe^2+^ to Fe^3+^, then autoclaved again. Oxidation of Fe^2+^ by air sparging was considered necessary to avoid abiotic oxygen consumption, as the digestate had high concentrations of Fe^2+^ originating from the Fe^3+^ used as precipitation chemicals in the primary wastewater treatment, and reduced to Fe^2+^ in the anaerobic digesters (Jonassen et al. 2022a). The sparging caused the pH to increase to 9.4 due the removal of CO_2_, requiring a final pH adjustment to 7.3 (with HCl). The same procedure was used for the field plot experiment, except that autoclaving was replaced by heat treatment: 70^0^C for 4 hours.

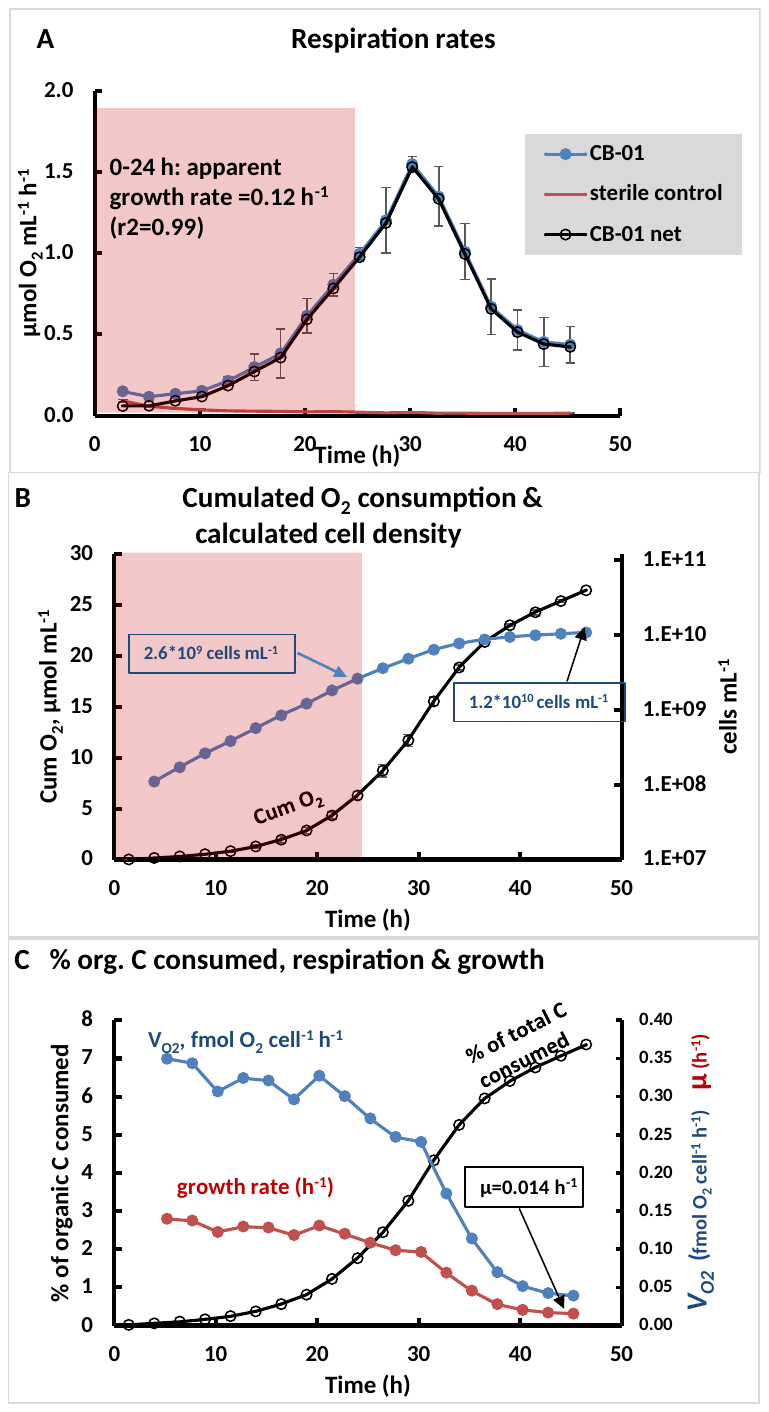
CB-01 was then grown aerobically in the the pretreated digestates, inoculated to an initial cell density of ~5*10^7^ cells mL^-1^, which were stirred and sparged with sterile air (filtered) at 23 ^0^C. To monitor the growth of CB-01, we transferred subsamples of each batch (after inoculation) to 120 mL vials (50 mL vial^-1^) with teflon coated magnetic stirring bars, which were placed in the incubation robot system for monitoring the O_2_ consumption (**Fig S7**).

**Fig S7**. **Cultivation of CB-01 in digestate.** The panels show the kinetics of O_2_-consumption as measured in the 50 mL subsamples placed in the incubation robot (4 replicate vials with CB-01, and 4 vials with sterile digestate).

Panel A shows the rates of O_2_ consumption in vials with CB-01 and in the sterile controls, and the net consumption by CB-01 (CB-01 minus sterile control). This increased exponentially during the first 24 hours, with apparent growth rate 0.12 h^-1^, which is much slower than in nutrient broth (0.29 h^-1^ **Fig S2**).

Panel B shows the cumulated O_2_ consumption by CB-01, and the estimated cell density assuming *Y*_O2_=4.06*10^14^ cells mol^-1^ O_2_ (**Fig S1**), reaching 1.2*10^10^ mL^-1^. The cell density quantified by qPCR for a similar experiment only 47% of the density based on O_2_ (**Table S2**), suggesting that *Y_O2_* for growth to high cell densities in digestate is ~50% of *Y_O2_* for optimal growth in nutrient broth (Fig S1).

Panel C shows estimated cell specific O_2_ consumption (*V_O2_*, fmol O_2_ cell^-1^ h^-1^), estimated growth rate, µ (h^-1^)= *V_O2_**Y, where Y = the measured growth yield by aerobic respiration (4.06 *10^14^ cells mol^-1^ O_2_), and the estimated fraction of organic C in the digestate (10 mg C mL^-1^) consumed by CB-01 (sum of CO_2_ and assimilated C).This suggests that ~1 % of the organic C in the digestate was easily available monomers, supporting rapid growth of CB-01 to a cell density of ~1*10^9^ cells mL^-1^ after 20 h, while subsequent growth was gradually declining as the organism utilized increasingly recalcitrant substrates, plausibly with a lower growth yield (Y_O2_).

**Table S2. Growth yield for CB-01 when growing to high cell densities.** The estimated growth of CB-01 in the digestate (Fig S7) was based on the growth yield as measured in batch cultures growing exponentially until limited by electron acceptors (4*10^10^ cells per mol O_2_ consumed, **Fig S1**), and the final cell density in these cultures was ≤ 2.7*10^9^ cells mL^-1^. The growth yield is plausibly lower for cultures that grow to higher cell densities, be it in nutrient broth or in digestate, due to a gradually declining growth rate induced by limitation of substrate supply. To assess the true growth yields under these conditions, we quantified the cell densities by real-time PCR (qPCR), and compared this with the estimated cell densities based on the oxygen consumption. The table shows this comparison both for growth to high cell densities in nutrient broth and digestate, confirming the hypothesis that growth yield is lower when the cultures reach high cell densities, and more so in digestate than in nutrient broth. Thus, the true cell density in the digestate used in the field experiments were approximately 50% of that estimated by the O_2_-consumption.

| Medium | Estimated cell density (10^9^ cells mL^-1^) | | qPCR-estimate as % of estimate based on O_2_-consumption |
| --- | --- | --- | --- |
|  | Based on O_2_ -consumption^*^ | Based on qPCR |  |
| Nutrient broth | 20.1 | 15.1 (stdev=0.7)** | 75 |
| Digestate | 10.5 | 4.92 (stdev=0.08)** | 47 |

* No standard deviation available for O_2_ consumption. The sample was taken from a single vial

** n=3 subsamples

**2C	Monitoring N_2_O emissions with a field robot**

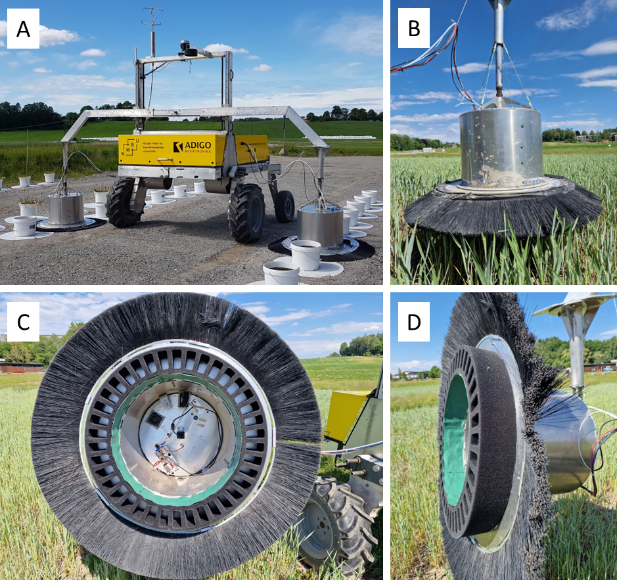
Emissions of N_2_O in all outdoor experiments were minitored by the “dynamic chamber” technique (Cowan et al. 2014, Hensen et al. 2006), operated by an autonomous Field Flux Robot (FFR) described byMolstad et al. (2014) and Molstad et al. (in prep.)

**Fig S8. The Field Flux Robot.** FFR (Panel A) operates two chambers (diameter 50 cm, height 50 cm), which are lowered over field buckets (Panel A) or onto the soil surface (Panel B). The chambers are equipped with cellular foam with gas tight flexible rubber coating which is compressed by deployment, and a circular brush skirting to function as a wind break (Panels C&D). The concentrations of N_2_O and CO_2_ in the two chambers are measured by circulating the chamber-air (intermittently for each chamber) via a Tunable Diode Laser N_2_O/CO-analyzer (DLT-100, Los Gatos Research, California, USA) and a CO_2_/H_2_O infrared gas analyzer (LI-840A, LI-COR Biosciences, Nebraska, USA), throughout a deployment time of 3 minutes.

**2D Field-bucket experiments**

Soils for the bucket experiments were collected from agricultural fields in southern Norway, spanning a range of soil characteristics:

**Table S2 Origin and characteristics of the soils**. The acid sandy silt soil (**S**) was taken from an agricultural field in Solør, Norway, dominated by fluvial sandy silt soils. The clay loam soils **L, I** and **N** were from different plots within a liming experiment near the Norwegian University of Life Sciences ((59°39’48.2”N 10°45’44.8”E), limed in 2014 (Nadeem et al. 2020), while **O** was a clay loam soil from the same area (hence with similar mineral components), but with a much higher content of organic C because it had been a peat-/wetland prior to cultivation. Soils **S, L, N** and **O** were used in the field bucket experiments. Soil **I** is the soil of the plots used for the field plot experiment.

| Soil: | pH (CaCl_2_) * | Tot C (%) ** | Tot N (%) ** | [NO_3_]  mg N kg^-1^ ** |
| --- | --- | --- | --- | --- |
| **S:** sandy silt soil | 4.15 | 0.75 | 0.07 | 30.2 |
| **L:** low-pH clay loam | 4.50 | 3.23 | 0.24 | 55.6 |
| **I**: intemediate pH clay loam | 6.13 | 3.23 | 0.24 | - |
| **N:** Neutral-pH clay loam | 6.70 | 3.21 | 0.25 | 21.1 |
| **O:** Organic-rich clay loam | 5.26 | 15.8 | 0.78 | 11.2 |

^1^ pH_CaCl2_ was measured after dispersing 10 g soil in 25 mL of 0.01 M CaCl_2_

^2^ Tot C= total organic C, Tot N = total organic N, and [NO_3_] = mg NO_3_N kg^-1^ soil dry weight.

Soil **L**, **I** and **N** were taken from different plots of a liming experiment on clay loam soil established and limed in 2014 (Nadeem et al., 2020). The low-pH clay loam (**L**) received no lime, the intermediate pH clay loam (**I**) was limed with 2.3 kg m^-2^ of dolomite, and the neutral-pH clay loam (**N**) was limed with 3 kg m^-2^ of finely ground calcite.

The soils used in the bucket experiments (**S**, **L**, **N** and **O**) were sieved (10 mm) in moist condition and mixed thoroughly before filling into the buckets. The conically shaped buckets (h = 21.5 cm, top diam. = 23.5 cm, bottom diam. = 21.5 cm) had a total volume of 8.6 L. A ~1 cm layer of gravel (4-8 mm diam.) was placed at the bottom, covered with a nylon fiber cloth to prevent eluviation of the soil by drainage. For soils **S**, **L** and **N**, 8 kg soil dry weight were filled into each bucket, packed by thumping the bucket on the ground till the soil had reached a bulk density of 1 kg L^-1^. For the organic rich clay loam soil, each bucket was filled with only 5.92 kg soil dry weight, reaching a bulk density of 0.74 kg L^-1^ after being packed to 8 L. The soil surface area of the buckets was 0.043 m^2^.

To secure equal initial amounts of NO_3_ m^-2^ for all soils, we mixed an amount of KNO_3_ to each soil to reach a level of 12 g N m^-2^ soil surface = 516 mg NO_3_-N bucket^-1^ (soil surface area = 0.043 m^2^). Digestate (480 mL bucket^-1^ = 11 L m^-2^ soil surface area) was mixed into the top ~10 cm of the soil by “harrowing”, using a small hand-held rake. We used autoclaved digestates in which CB-01 had been grown to ~6*10^9^ cells mL^-1^ , and as the control treatment we heat-treated this digestate (70 ^o^C, 2 h), which effectively killed the CB-01 cells (tested by measuring respiration, results not shown). As an additional control treatment, buckets received water only. The density of CB-01 cells per soil surface area immediately after application was 6.6*10^13^ cells m^-2^. The cell density in the upper 10 cm of the soil was ~6*10^8^ cells g^-1^ soil dry weight for the soils **S**, **L**, and **N** (bulk density = 1 kg L^-1^), and ~8*10^8^ g^-1^ for soil **O**.

The buckets were placed on 1 m^2^ plexiglass plates (1.5 mm), to avoid gas exchange with the soil below. The soil moisture (volumetric water content, m^3^/m^3^) and temperature (°C) in the upper 5.5 cm of the soil were monitored by four Teros 11 sensors, connected to an EM50 logger (Meter Group, Inc., WA, USA).

In the first experiment, using only Soil **N** (Table S2), starting 14.07.2021, rye gras was sown the day after the incorporation of the digestate, and the emissions were monitored for 90 days. Within this time span, we added 200 mL autoclaved and pH-adjusted digestate (4.6 L m^-2^) without CB-01 three times (after 19, 33 and 89 days), to induce transient bursts of N_2_O emission. By the end of each burst of N_2_O-emission induced by applying digestates, the upper 10 cm of the soil was sampled with an auger (diam. 1 cm) and stored in the freezer (-4 °C) until DNA extraction and subsequent molecular work. The auger was washed and sterilised with 70 % ethanol between each sampling.

In a follow up experiment, all soils were included and monitored for 10 days, with no re-fertilisation. Soil sampling was performed after the first peak of N_2_O emissions, as described for the 90 days bucket experiment.

The digestate application’s influence on soil pH was tested in the lab by mixing soil with the same type and amount of digestate as applied to the 0-10 cm soil layers of the field buckets (0.11 mL g^-1^ soil) ± 50 % to show the potential pH in pockets with higher or lower than average concentration of digestate. Water was added (if needed) together with digestate to reach the same water-filled pore space (WFPS, %) as in the field bucket experiment. The most prominent increase in soil pH was seen in the sandy silt soil, reflecting its low buffer capacity due to low content of clay and organic material (Table S1), both known to be crucial factors determining the soils’ buffer capacities (Curtin et al. 1996).

**Table S3. Soil pH changes due to application of digestate.** The table shows pH(CaCl_2_) in the four soils as affected by applying 0.055, 0.11 and 0.165 g digestate g^-1^ soil, which is 50, 100 and 150 % of the amounts added to the soils in the field bucket experiment (0.11 mL digestate g^-1^ soil). Water was added (if needed) to reach a WFPS (%) equivalent to that in the buckets after digestate application (last column).

| **Soil type** | **no digestate** | **0.055 mL digestate g^-1^ soil** | **0.11 mL**  **digestate g^-1^ soil** | **0.165 mL digestate g^-1^ soil** | **WFPS (%)** |
| --- | --- | --- | --- | --- | --- |
| **S:** Sandy silt soil | 4.15 | 5.02 | **5.33** | 5.59 | 49 |
| **L:** Low-pH clay loam | 4.50 | 4.87 | **5.00** | 5.11 | 53 |
| **N:** Neutral-pH clay loam | 6.70 | 6.75 | **6.75** | 6.74 | 54 |
| **O:** Organic-rich clay loam | 5.26 | 5.42 | **5.52** | 5.59 | 45 |

**2E Field plot experiments**

To assess the effect of CB-01 in digestate on soil N_2_O emission under field conditions, we established small (0.5 m^2^) test plots within larger field plots (8 m X 3 m) of a soil liming experiment (limed in 2014) on clay loam soil (Nadeem et al. 2020, Byers et al. 2021) and re-limed with 174 g dolomite m^-2^ in 2019. We used the plots with soil I (Table S2) that were previously limed with dolomite to pH(CaCl_2_) = 6.13 (stdev = 0.10), and within each of the 6 replicate plots, we established two 0.7 m X 0.7 m test plots side by side (distance = 30 cm), fertilized with autoclaved digestate in which CB-01 had been grown to a cell density of ~6*10^9^ cells mL^-1^. We applied 4.5 L digestate per plot (= 9 L m^-2^), which was mixed into the upper ~10 cm of the soil by a hand-held cultivator. The intital density of CB-01 was 5.4*10^13^ cells m^-2^. If distributed throughout the the soil layer that was sampled for analyses (0-10 cm depth = 125 kg soil dry weight m^-2^, assuming a bulk density of 1.25 kg L^-1^), the initial cell density in the soil would be 4.3*10^8^ cells g^-1^ soil. Soil samples for determining CB-01 abundance were taken from each plot (3 replicate samples) before incorporation of digestate with CB-01, 9 days later, and after 10 months. The soil samples were stored in the freezer (-20 °C) until DNA extraction and following quantification by PCR.

The 0.5 m^2^ test plots were situated along the boardwalk for the autonomous field flux robot (FFR), which was used to monitor the N_2_O emission.

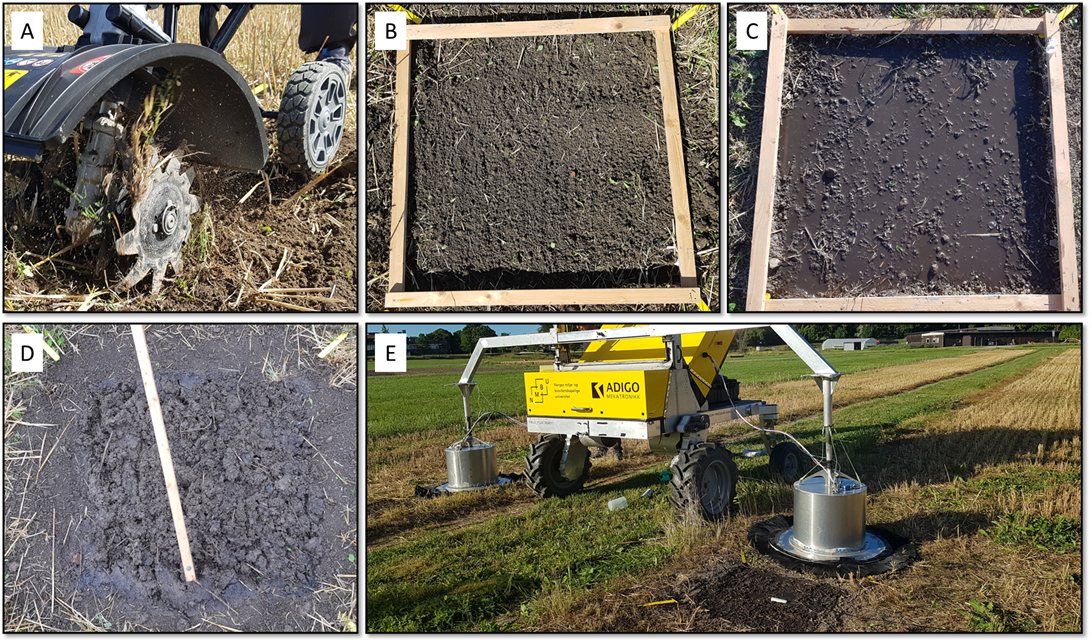

**Fig S9. Field plot experiments.** The top ~10 cm of the soil was first loosened by a cultivator (Panel A). After evening out the surface with a rake, we placed a 0.7 X 0.7 m frame onto the surface (Panel B), and poured digestate onto the surface within the frame (Panel C). One day later the top soil (0-10 cm) was harrowed by a hand held tool (Panel D), and N_2_O emissions were monitored by the robot (Panel E).

**2F Calculations of emissions and statistical analyses**

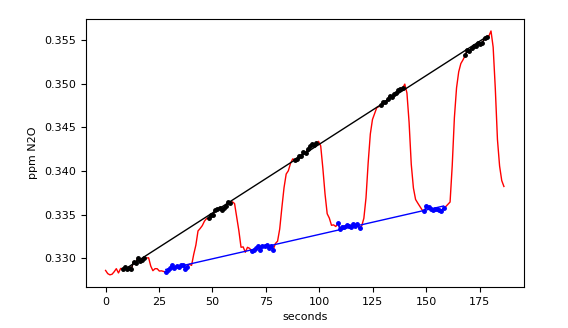

**Fig S10 Recorded signal of N_2_O concentration from a single measurement with the FFR.** The jagged shape of the curve is due due to valves in the robot switching every 20 seconds, alternating circulating air from the left and right chamber through the laser instrument. The two straight lines are regression lines, the slopes of which is used to calculate the emission fluxes. The difference in the steepness of the regression lines is caused by a difference between the fluxes from the soils the chambers have been lowered onto.

Emissions: From the slope of the N_2_O regression lines the flux of N_2_O is calculated by the equation

$q_{N2O} = \frac{{10}^{-6} a h p}{R T}$,

where *q_N2O_* is the flux of N_2_O (mol m^-2^ s^-1^), *a* is the slope of the regression line (ppm s^-1^), *h* is the height (i.e., the volume divided by the ground surface area) of the chamber (m), *p* is the pressure (Pa), *R* is the universal gas constant (J mol^-1^ K^-1^) and *T* is the temperature (K).

For graphic presentation of the emissions, we used the Gausian Kernel smoother (Hastie et al 2009) to plot floating averages for each treatment (continous curves) together with individual measurements (as dots); Figs 2,4,5 in main paper.

Cumulative N_2_O emissions over a period of time are approximated by using the trapezoidal rule on the estimated fluxes. ( $\int q_{N2O}(t) dt\approx\sum(q_{N2O}\left( t_{i} \right)+q_{N2O}\left( t_{i+1} \right))(t_{i+1}-t_{i})/2$ ). This was done for each individual bucket and field plot.

The field plot experiment yielded paired data – six pairs (*X_i,_ Y_i_*)*, i* = 1 .. 6, where *X_i_* are cumulative emissions from plots treated with NNRB, and Y_i_ are cumulative emissions for control plots. This gives six ratios *R_i_* = *X_i_*/*Y_i_* . Confidence intervals for the mean of the ratios, 1/6 Σ*R_i_* , for two time periods were made with a Student’s t distribution (assuming that the ratios were normally distributed.). These confidence were similar to confidence intervals found by the Fieller method for ratios of paired data and also by simple nonparametric bootstrapping (Efron and Tibshirani, 1994).

Since the field bucket experiments did not yield paired data, flux reduction statistics are calculated as ratios of means, rather than means of ratios, of cumulative fluxes. Confidence intervals of these ratios were made by the Fieller method for unpaired data (Motulsky, 1995, p 285) and by simple nonparametric bootstrapping (the results were similar). The 95% coverage of the Fieller confidence intervals were tested by numerical simulations and a bootstrap-calibration of the confidence level was made, with negligible effects on the confidence intervals.

**2G	Tracing CB-01 in digestate and soil**

To measure growth of CB-01 in digestate, and its survival in soil, we used quantitative PCR (qPCR), with primers that are specific to *Cloacibacterium* strains, developed by Allen et al. (2006). The primers target the complementary parts of the following sequence of the 16S rRNA gene: 5’-TATTGTTTCTTCGGAAATGA (Cloac-001f) and 5’-ATGGCAGTTCTATCGTTAAGC (Cloac-001r).

DNA was extracted with the DNeasy PowerSoil Pro Kit (Qiagen) according to the manufacturers protocol, except for the first step: Bead beating of the cells was done at 4.5 m s-1 for 45 s in a FastPrep-24™ (MP Biomedicals, LLC, CA, USA), instead of a vortex. To measure the concentration of DNA in the extract, we used a Qubit dsDNA HS Assay Kit (Thermo Fisher Scientific, USA). The number of CB-01 16S gene copies in extracted DNA was quantified using a CFX96 Touch™ Real-Time PCR Detection System (Bio-Rad, USA), running for 15 min at 95 °C followed by 40 cycles of denaturation (30 s at 95 °C), annealing (30 s at 55 °C) and elongation (45 s at 72 °C). The master mix contained 0.2 µM of each primer (Cloac-001f and Cloac-001r), and 1x HOT FIREPol® EvaGreen® qPCR Supermix (Solis BioDyne).

For calibration, we used DNA extracted suspensions of washed cells containing 10^3^, 10^4^, 10^5^, 10^6^, 10^7^ and 10^8^ cells mL^-1^, resulting in 2.4*10^1^-2.4*10^6^ 16S templates per PCR tube (taking dilution into account, and the fact that each genome of CB-01 contains three 16S gene). To enable the use of the Cq values to estimate copy numbers, we used Generalized Reduced Gradient Solver in Excel to fit the model (equation 1) to the data:

$$N= \frac{N_{T}}{{(2\cdot e)}^{Cq}} \left( 1 \right)$$

where ***N*** is the initial number of 16S templates in the PCR tube, ***N_T_*** is the number of amplicons per tube needed for signal detection (above background), ***e*** is the efficiency of the PCR amplification, and ***Cq*** is the number of cycles needed for detection of a signal. The fitted parameters were ***N_T_* = 7.68*10^10^ copies per tube and *e* = 0.85 (85 % efficiency)**.

An independent dataset was provided by running qPCR with the same primers on extracted DNA from suspensions of unwashed CB-01 cells (in nutrient broth) with densities 10^4^, 10^5^, 10^6^, 10^7^ and 10^8^ cells mL^-1^. The log_10_ values of cell densities estimated by the Cq values were on average 104 % of the expected value, with a standard deviation of 6 %.

When using qPCR to estimate the CB-01 abundance in soil and digestate, inhibition of the polymerase can result in too high Cq numbers, hence resulting in underestimation of the gene abundance (Lim et al. 2016). To investigate this, we spiked the different soils and the digestate with 10^9^ CB-01 cells g^-1^ soil dry weight and mL^-1^ digestate, respectively, extracted DNA from 0.2 g soil and 0.2 mL digestate, eluted to a 50 µL DNA solution for each material, which was then diluted in 10-fold steps from 0 (undiluted) down to 1/10^7^. The results (**Figure S10)** show a reasonable fit between model (predicted) and measured Cq values for all materials if diluting the extracted DNA to ≤ 1/10, except for the clay loam pH 6.13 soil, which required dilution to ≤1/100 to eliminate inhibition.

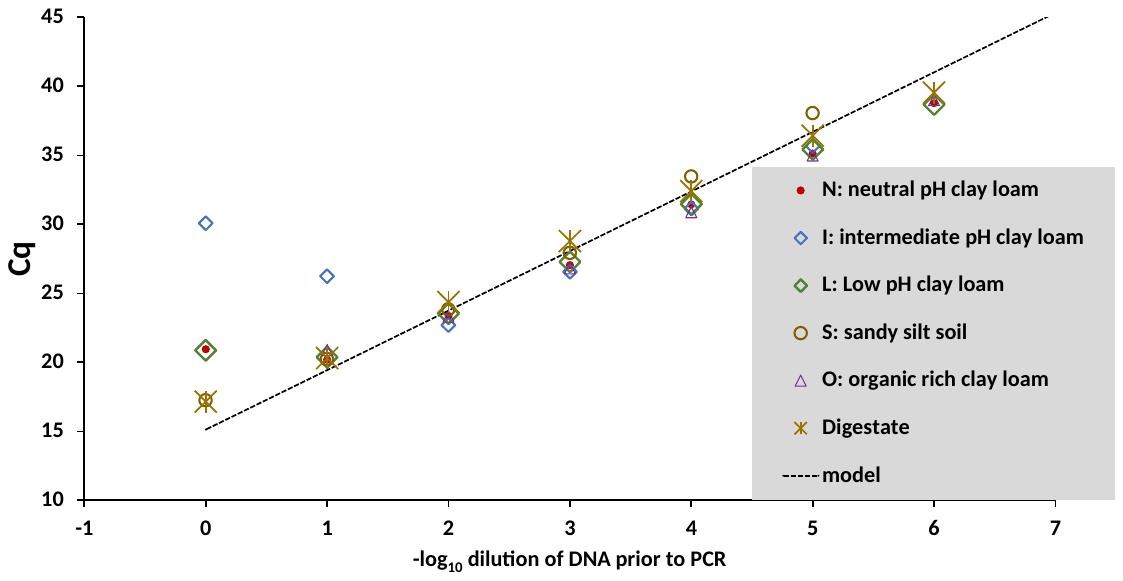
**Fig S11 Inhibition of qPCR.** Soils and digestate was spiked with CB-01 cells (10^9^ cells g^-1^ soil dw and mL^-1^ digestate, respectively). DNA was extracted (0.2 g soil and 0.2 mL digestate was extracted, eluted in a 50 µL volume. From undiluted, and 10-fold dilutions of this, 2µL were transferred to PCR tubes for amplification of CB-01 16S genes. The plot shows the Cq values plotted against dilution, together with the model *N = N_T_/(2*e)^Cq^* as parameterized (see text): ***N_T_***=7.68*10^10^ copies per tube and *e*=0.

The result can be used to approximate the lower limit for detection of CB-01 in soils and digestate: A cautious upper limit for Cq values to be trusted is 40, i.e. 34 templates per PCR tube (equation 1). The polymerases were evidently inhibited by using undiluted DNA in the reaction (**Fig S10**), hence a 1/10 dilution of the extracted DNA is needed for all soils except soil **I**, for which 1/100 dilution is required. This means that the PCR tube can maximally be loaded with DNA from 0.8 mg soil (0.08 mg for soil **I**) and 0.8 µL digestate. This implies a limit of detection around 4.3*10^4^ templates g^-1^ soil (4.3*10^5^ for soil I due to dilution to 1/100) and mL^-1^ digestate, or 1.4*10^4^ CB-01-genomes g^-1^ soil and mL^-1^ digestate (since the genome contains 3 copies of the 16S gene).

The real limit of detection for a CB-01 inoculum in soil and digestate could be higher than this, if indigenous genes are amplified with the primers. This was tested by running PCR on soil and digestates which had not been spiked with CB-01, along with analyzing spiked samples in various experiments. The results are summarized in Table S4. Since there were several tubes with a negative result (Cq>40), average values cannot be calculated. A cautious judgment would be that the “background” PCR signal of the soil is Cq=39-38, which is equivalent to 67-107 templates per PCR tube, or 21-36 CB-01-genomes per tube. For all soils except **I**, we used the Cq values for the PCR tubes loaded with 1/10 dilutions, which were thus loaded with DNA from 0.8 mg soil. For these, the background PCR signal is equivalent to 2.6-4.8*10^4^ CB-01 genomes g^-1^, while 10 times higher for soil **I** (due to 1/100 dilution of the DNA from this soil). For digestate, the average Cq was 31.98 (Table S4), which means that the untreated digestate contains 3.2*10^6^ CB-01 16S-templates mL^-1^, or 1.1*10^6^ CB-01 genomes mL^-1^.

**Table S4 PCR-results for soils and digestate without spiking with CB-01**. The table shows the total number of samples analyzed for each soil and for the digestate, the number of samples with Cq>40 (estimate not available), and the average for those <40. The results suggest that the background Cq values are >38, which implies that the background is <1.25*10^5^ 16S-templates g^-1^, hence <4.2*10^4^ CB-01 genomes g^-1^ soil. The background value of digestate is a bit higher, with a Cq of 31.98, corresponding to about 3.2*10^6^ 16S-templates mL^-1^, or 1.1*10^6^ CB-01 genomes mL^-1^ digestate.

| Material | Total number of samples | Number of samples with Cq<40 | Average Cq for samples with Cq<40 |
| --- | --- | --- | --- |
| **N:** Neutral-pH clay loam | 8 | 2 | 39.33 |
| **I:** Intermediate-pH clay loam | 12 | 7 | 39.48 |
| **L:** Low-pH clay loam | 3 | 1 | 37.78 |
| **S:** Sandy silt soil | 3 | 0 | - |
| **O:** Organic-rich clay loam | 3 | 2 | 39.8 |
| Digestate | 3 | 3 | 31.98 |

**2H** **Survival of CB-01 in soil, laboratory incubation**

A soil incubation experiment was designed to assess the survival of CB-01 in soil, vectored by digestate, under constant temperature and moisture conditions, and without any subsequent incorporation of digestate (thus contrasting the field bucket experiment, Fig 2 in main paper). CB-01 was first grown to ~6*10^9^ cells mL^-1^ in autoclaved, aerated and pH-adjusted digestate (as for the field experiments). Neutral-pH clay loam soil (soil N **Table S2**) was portioned into a set of 50-mL Falcon tubes (9.4 g soil dry weight, moisture content = 0.5 mL g^-1^ soil dry weight). To each tube, 4.2 mL sterile water and 0.85 g digestate (with CB-01) was dripped onto the soil. The tubes were stored in a dark moist chamber at 15 ^0^C, with loose lids to allow exchange of air. Control tubes received only sterile water. At intervals, 2 replicate tubes were frozen (-20 ^0^C) for quantification of CB-01 gene abundance by qPCR as described above.

**2I Extrapolating to national emission reductions**

We use the emissions quantified with the GAINS model (Winiwarter et al. 2018, Amann et al. 2011) for 2030 in Europe to estimate the possible reductions of the measure.

The experiments described in this paper demonstrate marked emission reductions on all soils tested, over extended periods. The strongest reductions have been seen for the initial N_2_O peak immediately after fertilization, but NNRB has shown to remain active over a period of 90 days. Cumulative emissions over the whole period have been reduced by at least 41% (for clay loam soils), up to 95% reduction. We may disregard the case of smallest reduction as also the emissions from these soils are rather small, but the organic loam soils (55% reductions) need to be considered. Consistent with the uniform emission factor used in GAINS (from IPCC, 2006) of 1% of N applied to be emitted as N_2_O for all conditions of crops, soil or type of fertilizer added, we also choose to apply a uniform reduction factor of 60% of emission reductions due to NNRB which we consider a conservative estimate. In Table S5, emission reductions are shown by EU country for 2030 if emissions from application of liquid manure only is reduced by 60%. This assumption is based on the understanding that liquid manure can easily be treated in biodigesters. Höglund et al. (2020) assume, for purpose of methane abatement, anaerobic digestion becomes profitable only for large agricultural entities of at least 100 livestock units. According to GAINS numbers, this concerns 70% of all farms in the EU, which more probably reflect liquid than solid manure systems, so the above estimate remains valid for the major fraction of liquid manure available. Indirect emissions as well as other soil emissions due to grazing, mineral fertilizer additions or application of farmyard manure (solid manure systems) have been left unchanged. Note that the GAINS model (in agreement with IPCC, 2006) does not account for potentially increased emissions due to dry periods or freeze-thaw cycles (the latter considered to potentially contribute as much as 17-28% to global soil emissions: Wagner Riddle et al. 2017) while covering increased emissions from cropping histosols.

Under these assumptions, total N_2_O emissions from Europe decrease by 2.7% due to NNRB introduced. This figure is higher in countries that have a high share of liquid manure systems in their agriculture, hence for EU27 (27 EU member countries) the corresponding figure is 4.0%, if NNRB were used for all manure nitrogen applied from liquid manure systems.

If it were possible to extend the NNRB-technology, using solid manure and plant residues as substrates and vectors, we speculate emission reductions could be achieved for all mineral and natural fertilizer actively applied on fields. Ongoing work has shown that while *Clocaibacter* CB-01 grows to high cell densities in plant residues, new strains which grow in manure have been enriched and isolated (unpublished results). Although further development will be needed to implement this , it is relevant to estimate their impacts. Applying NNRB also to these other substrates at the same reduction efficiency could decrease European emissions as well as EU27 emissions by about a quarter (24% and 23%, respectively). For agricultural emissions only, this means that roughly a third (31%) could be eliminated.

It needs to be pointed out that an emission reduction of 60% as derived here for NRB is much larger than emission reductions typically reported for N_2_O abatement measures. E.g., GAINS assumes nitrification inhibitors to be able to reduce emissions by as much as 38%, and high-tech mechanical fertilizer saving technologies (“variable rate application”) to be able to save 24% of the emissions only (Winiwarter et al. 2018). Of note, the percent reduction of N_2_O-emission by the NNRB-technology is plausibly unaffected by “variable rate application” and nitrification inhibitor, since the target for NNRB is to reduce the N2O/N2 product ratio of denitrification, while the two others target the concentration of NO_3_^-^ and nitrification, respectively.

**Table S5: Potential emission reductions as a consequence of implementing NNRB measures** (anthropogenic emissions in kt N_2_O per year projected for 2030)

|  | **Total N_2_O emissions**  **Kt N_2_O y^-1^** | **Reduction by NNRB**  **kt N_2_O y^-1^** | | **% Reduction of total emissions** | | **% reduction of agricultural emissions** |
| --- | --- | --- | --- | --- | --- | --- |
|  |  | NNRB in liquid manure only | NNRB with all N- fertilizers | NNRB in liquid manure only | NNRB with all N- fertilizers | NNRB with all N- fertilizers |
| Albania | 4.3 | 0.10 | 0.85 | 2.3% | 20% | 24% |
| Austria | 13.4 | 0.80 | 2.99 | 5.9% | 22% | 34% |
| Belarus | 44.7 | 0.20 | 8.29 | 0.4% | 19% | 23% |
| Belgium | 23.0 | 0.96 | 5.11 | 4.2% | 22% | 33% |
| Bosnia-Herzegovina | 3.9 | 0.12 | 0.75 | 3.0% | 19% | 25% |
| Bulgaria | 13.6 | 0.09 | 4.44 | 0.6% | 33% | 40% |
| Croatia | 6.0 | 0.12 | 1.67 | 2.0% | 28% | 34% |
| Cyprus | 0.9 | 0.05 | 0.15 | 5.9% | 16% | 21% |
| Czech Republic | 19.2 | 0.18 | 5.11 | 0.9% | 27% | 37% |
| Denmark | 18.8 | 1.83 | 5.24 | 9.8% | 28% | 33% |
| Estonia | 3.2 | 0.06 | 0.73 | 1.8% | 23% | 31% |
| Finland | 17.6 | 0.45 | 2.93 | 2.5% | 17% | 25% |
| France | 143.3 | 5.16 | 36.76 | 3.6% | 26% | 31% |
| Germany | 132.5 | 7.46 | 30.09 | 5.6% | 23% | 31% |
| Greece | 14.0 | 0.18 | 2.60 | 1.3% | 19% | 25% |
| Hungary | 17.4 | 0.18 | 5.85 | 1.1% | 34% | 41% |
| Iceland | 1.0 | 0.04 | 0.24 | 4.3% | 24% | 28% |
| Ireland | 30.6 | 1.40 | 5.73 | 4.6% | 19% | 20% |
| Italy | 60.0 | 3.08 | 13.28 | 5.1% | 22% | 33% |
| Kosovo | 1.2 | 0.02 | 0.18 | 1.5% | 14% | 28% |
| Latvia | 4.9 | 0.08 | 0.86 | 1.6% | 17% | 20% |
| Lithuania | 13.2 | 0.30 | 2.58 | 2.2% | 20% | 22% |
| Luxembourg | 1.2 | 0.05 | 0.18 | 4.3% | 15% | 26% |
| Malta | 0.2 | 0.01 | 0.04 | 2.3% | 17% | 32% |
| Moldavia | 2.6 | 0.03 | 0.58 | 1.0% | 22% | 28% |
| Montenegro | 0.5 | 0.01 | 0.06 | 1.5% | 11% | 19% |
| Netherlands | 36.0 | 2.69 | 5.28 | 7.5% | 15% | 23% |
| Northern Macedonia | 1.9 | 0.04 | 0.29 | 2.0% | 15% | 25% |
| Norway | 11.1 | 0.41 | 2.17 | 3.7% | 20% | 30% |
| Poland | 81.3 | 2.32 | 20.65 | 2.8% | 25% | 33% |
| Portugal | 11.4 | 0.32 | 1.91 | 2.8% | 17% | 23% |
| Romania | 28.8 | 0.48 | 8.00 | 1.7% | 28% | 35% |
| Russia | 291.4 | 2.22 | 79.37 | 0.8% | 27% | 33% |
| Serbia | 10.7 | 0.31 | 2.78 | 2.9% | 26% | 37% |
| Slovakia | 7.3 | 0.08 | 2.09 | 1.1% | 29% | 40% |
| Slovenia | 2.4 | 0.16 | 0.46 | 6.8% | 19% | 31% |
| Spain | 62.1 | 2.47 | 14.75 | 4.0% | 24% | 32% |
| Sweden | 18.7 | 0.49 | 3.47 | 2.6% | 19% | 29% |
| Switzerland | 9.3 | 0.57 | 2.01 | 6.1% | 22% | 31% |
| Turkey | 109.4 | 0.81 | 23.45 | 0.7% | 21% | 28% |
| Ukraine | 73.4 | 0.34 | 18.61 | 0.5% | 25% | 34% |
| United Kingdom | 89.4 | 1.69 | 19.39 | 1.9% | 22% | 29% |
| All Europe (incl. non-European parts of Russia and Turkey) | 1435.9 | 38.51 | 341.97 | 2.7% | 24% | 31% |
| sum EU27 | 781.0 | 31.46 | 182.97 | 4.0% | 23% | 31% |

**3	Effect of CB-01 on the soil microbiome**

Microbial community composition was examined by amplicon sequencing of the 16S rRNA gene V3-V4 region. Purified DNA from soil samples was sent to Novogene Europe for amplification, library preparation and sequencing to generate 250 bp paired-end reads using the Illumina Novoseq platform. Reads, after primer removal, were processed using GHAP (V2.4), an in-house amplicon clustering and classification pipeline built around Usearch (V11.0.66) (Edgar, 2010), the RDP classifier (V2.13) (Wang et al. 2007) and locally written tools for generating OTU tables. Reads were processed using default quality control and trimming parameters. Clustering was performed at both 97% and 100% similarity to generate OTUs and zOTUs, respectively. The 16S rRNA gene sequence of *Cloacibacterium* sp. CB-01 (GCA_907163125) was then matched against the OTU and zOTU representative sequences using the Usearch usearch_global command at 97% similarity and 99% similarity, respectively, to determine which OTU and zOTUs circumscribe the *Cloacibacterium* sp. CB-01 inoculant. From visual inspection it appeared that two zOTUs (zotu45 and zotu611) may circumscribe *Cloacibacterium* sp. CB-01 due to shared abundance profiles and taxonomic classifications. To confirm that these two zOTUs both matched to *Cloacibacterium* sp. CB-01 the two representative sequences were BLAST searched (Alstschul et al, 1990) against the *Cloacibacterium* sp. CB-01 genome, where it was observed that both zOTU sequences matched closely to two separate regions of the genome, presumably harbouring multiple slightly divergent copies of the 16S rRNA gene. To confirm, the two 16S rRNA genes from the *Cloacibacterium* sp. CB-01 genome were matched back against the zOTU representative sequences using the usearch_global command at 99% similarity where they matched to both zotu45 and zotu611, separately. Due to this zotu45 and zotu611 were combined for downstream analyses.

To assess the impact of the various treatments on the soil microbial communities alpha and beta diversity measures were calculated for microbial communities from all samples using the OTU tables generated above. OTU tables were first modified by removing the OTU circumscribing *Cloacibacterium* sp. CB-01 (OTU_27) before rarifying the tables to 72 846 reads per sample using the Usearch otutab_rare command. Shannon’s (Shannon, 1948) and Simpson’s (Simpson, 1949) diversity indices were calculated using the Usearch -alpha_div command and beta diversity measures calculated using the Usearch -beta_div command. Jaccard’s dissimilarity measures (Jaccard, 1912) were then used to generate MDS plots using the Scikitlearn MDS module (Pedregosa, et al. 2011).

The beta diversity as shown by Jaccard’s dissimilarity measures indicated that early during the soil incubation period there is greater between sample variation both within treatments as well as between soils treated with live CB-01 and those treated with water or dead CB-01. Indicating an effect of CB-01 on the soil microbial communities (**Fig S11**). This effect, however, disappears by the final time point where samples from live CB-01, dead CB-01 and water treated soils cluster together, suggesting the effect of live CB-01 on native soil microbial communities is transient and microbial soil communities are not affected in the longer term by live CB-01 addition. It should be noted that the effect over time throughout the experiment is also a much larger source of microbial community variation than the addition of live CB-01 cells. Presumably due to disturbances to the soil from digging, sieving and packing of pots. Similarly, no systematic effects are observed on the alpha diversity of soil microbial communities throughout the experiment indicating that the CB-01 treatment does not reduce the complexity or evenness of soil microbial communities when added to soils with digestate organic matter as can be seen in the Shannon’s and Simpson’s diversity measures of samples taken throughout the experiment (**Fig S12** and **S13**).

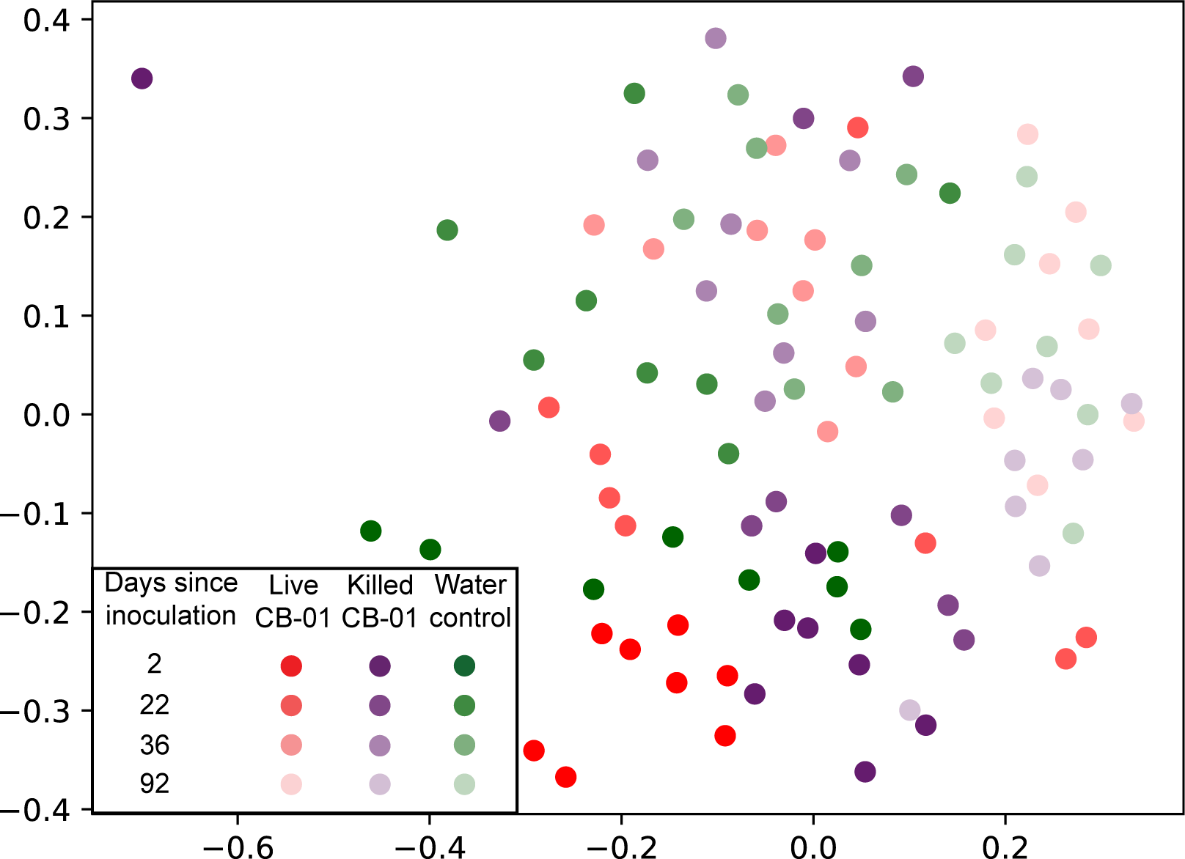

**Fig S12 MDS plot based on community composition of soils inoculated with CB-01.** This panel shows an MDS plot based on Jaccard’s dissimilarity measures for microbial communities in the soil of the field bucket experiment treated with live CB-01 in digestate, killed CB01 in digestate and water treatment only.

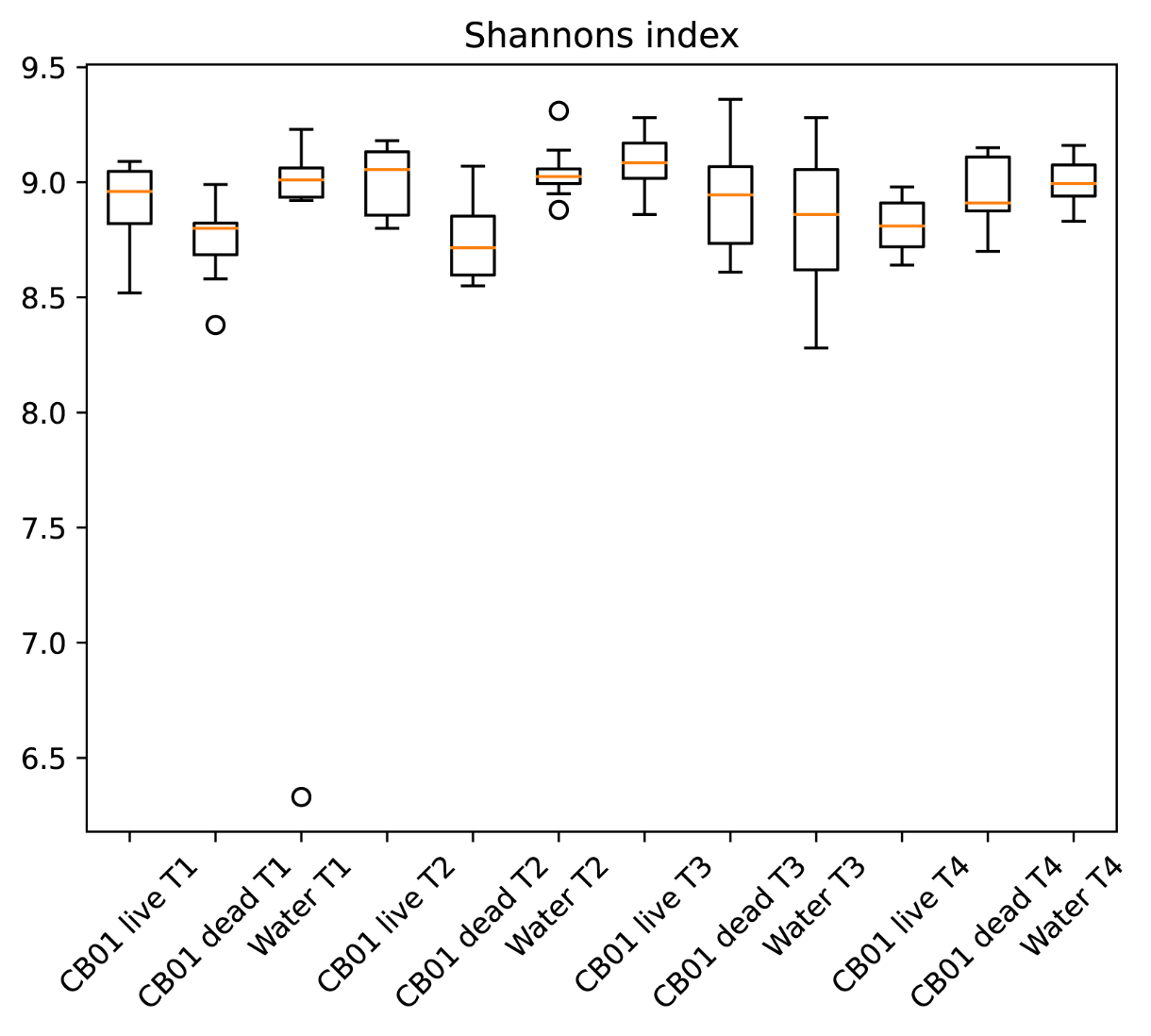

**Fig S12 Box and whisker plot showing the Shannon’s diversity indices for soil microbial communities.** This panel shows the Shannon’s diversity indices for microbial soil communities sampled throughout the field bucket experiment treated with live CB-01 in digestate, killed CB-01 in digestate and water treatment only.

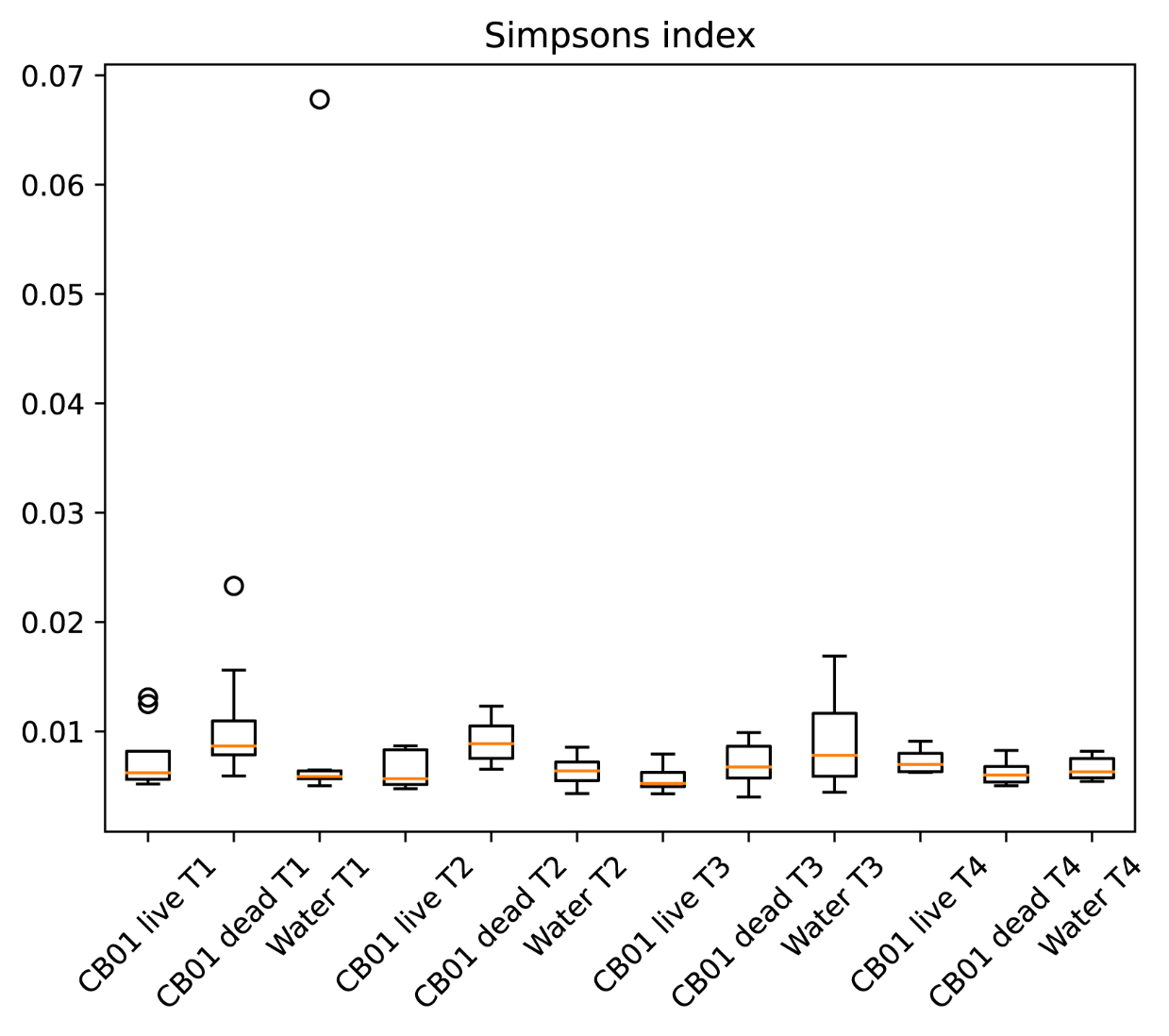

**Fig S13 Box and whisker plot showing the Simpson’s diversity indices for soil microbial communities.** This panel shows the Simpson’s diversity indices for microbial soil communities sampled throughout the field bucket experiment treated with live CB-01 in digestate, killed CB-01 in digestate and water treatment only.

**4	Survival of CB-01 in the field plot experiment**

Table S6 and Figure S14 show the estimated average abundance of CB-01 genomes (unit= 1000 g^-1^ soil dry-weight) in the paired field plots (a-f), i.e. adjacent plots inocculated with live and heat-killed Cloacibacter. The number of cells added with the digestate was 4.3*10^9^ g^-1^ soil dryweight. The genome abundance in the soil samples taken 9 days after fertilization are remarkably variable, suggesting a fast initial decline in two of the plots with live CB-01 (plot c&e), at rates comparable to the rapid initial decline in the laboratory incubation experiment (Figure 6 in the main paper), tentatively ascribed to protozoal grazing. The variability suggests a patchy distribution of protozoa within the field.

Of note, the genome abundance after 280 days in the plots with live CB01 are in some agreement with the first order decline rates estimated for the field bucket experiment: The decline in genome abundance from 4.3*10^9^ at time 0 to 1.5*10^6^ at time 280 days implies an average first order decay rate of 0.028 d^-1^, or an average half life = 25 days.

**Table S6. Estimated abundance of CB-01 genomes in the field plot experiment,** 9 and 280 days after fertilization with CB-01 containing digestate (live or heat killed). The numbers are the average genome abundance based on qPCR anallysis of 3 replicate samples for each indivdual plot. Unitis thousand genomes g^-1^ soil dry-weight. Initial numbers of CB-01 (at time 0) was ~6*10^9^ cells g^-1^ soil dry-weight, based on analysis of the digestate (Fig S7).

|  | **Plots with live CB-01** | | | | **plots with heat-killed CB-01** | | | |
| --- | --- | --- | --- | --- | --- | --- | --- | --- |
|  | **Time 9 days** | | **time=280 days** | | **Time 9 days** | | **time=280 days** | |
| **plot-pair** | **avg** | **se** | **avg** | **se** | **avg** | **se** | **avg** | **se** |
| average | 698 007 | 194 735 | 2 331 | 1 891 | 7 064 | 3 446 | 70 | 33 |
| b | Na* | - | 3 510 | 1 321 | 98 195 | 90 261 | 203 | 17 |
| c | 1 531 | 533 | 652 | 469 | 424 | 196 | 160 | 79 |
| d | 471 431 | 189 461 | 1 229 | 650 | 58 446 | 29 613 | 176 | 14 |
| e | 2 298 | 1 619 | 814 | 531 | 43 173 | 27 419 | 81 | 44 |
| f | 608 617 | 599 881 | 677 | 447 | 599 | 281 | 191 | 82 |
| **average** | **356 377** | **149 141** | **1 536** | **471** | **34 650** | **16 092** | **147** | **23** |

*samples lost

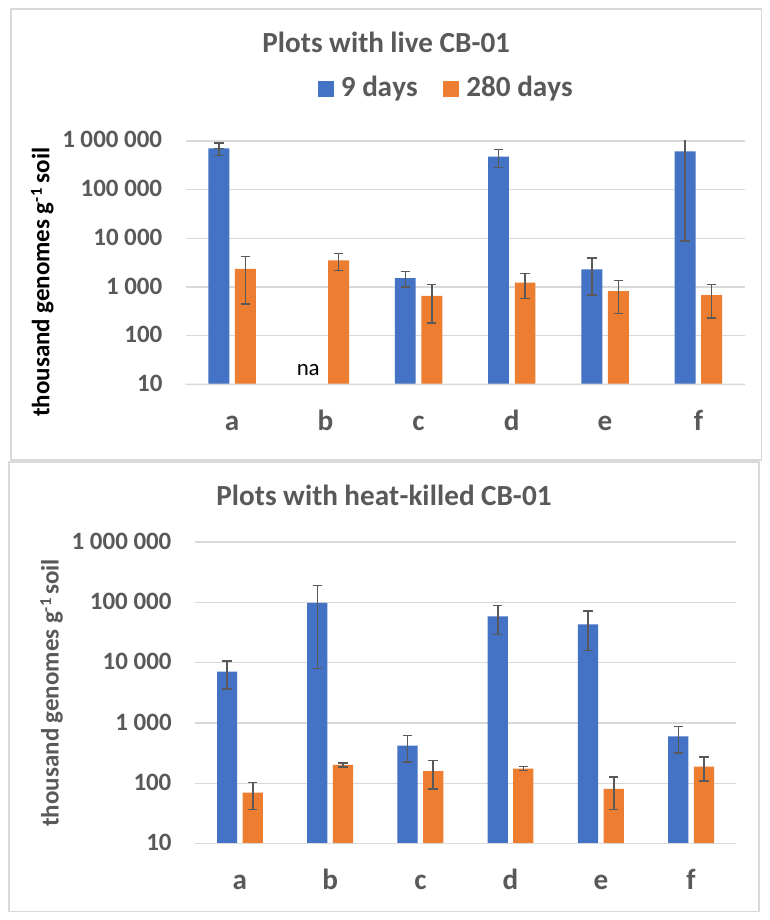

**Fig S14 Graphic presentation of the data in Tale S6**. Standard error shown as vertical lines (n=3).
